# Dendritic branch structure compartmentalizes voltage-dependent calcium influx in cortical layer 2/3 pyramidal cells

**DOI:** 10.1101/2022.01.11.475883

**Authors:** Andrew T. Landau, Pojeong Park, J. David Wong-Campos, He Tian, Adam E. Cohen, Bernardo L. Sabatini

## Abstract

Back-propagating action potentials (bAPs) regulate synaptic plasticity by evoking voltage-dependent calcium influx throughout dendrites. Attenuation of bAP amplitude in distal dendritic compartments alters plasticity in a location-specific manner by reducing bAP-dependent calcium influx. However, it is not known if neurons exhibit branch-specific variability in bAP-dependent calcium signals, independent of distance-dependent attenuation. Here, we reveal that bAPs fail to evoke calcium influx through voltage-gated calcium channels (VGCCs) in a specific population of dendritic branches in cortical layer 2/3 pyramidal cells, despite evoking substantial VGCC-mediated calcium influx in sister branches. These branches contain VGCCs and successfully propagate bAPs in the absence of synaptic input; nevertheless, they fail to exhibit bAP-evoked calcium influx due to a branch-specific reduction in bAP amplitude. We demonstrate that these branches have more elaborate branch structure compared to sister branches, which causes a local reduction in electrotonic impedance and bAP amplitude. Finally, we show that bAPs still amplify synaptically-mediated calcium influx in these branches because of differences in the voltage-dependence and kinetics of VGCCs and NMDA-type glutamate receptors. Branch-specific compartmentalization of bAP-dependent calcium signals may provide a mechanism for neurons to diversify synaptic tuning across the dendritic tree.

## Introduction

The synaptic tuning properties of neurons are regulated by voltage-dependent calcium influx, which is typically evoked by back-propagating action potentials (bAPs) (Magee and Johnston, 1997, Sjöström, 2002, Hausser and Mel, 2003). Many investigations have focused on timing-dependent plasticity rules, such as spike-timing dependent plasticity (STDP), in which the delay between synaptic input and the bAP determines if an induction protocol results in long-term potentiation (LTP), depression (LTD), or maintenance of synapse strength (Feldman, 2012). However, other studies have shown that the outcome of plasticity induction protocols also depends on the synaptic location within the dendritic tree (Golding et al., 2002, Froemke et al., 2005, Sjostrom and Hausser, 2006, Gordon et al., 2006). The same protocol can evoke a different sign, magnitude, or temporal profile of synaptic plasticity if applied to synapses in proximal or distal dendritic compartments. Such location-dependent plasticity rules may bolster the integrative capacity of individual neurons by supporting divergent tuning properties in basal and apical dendritic arbors (Larkum et al., 1999, Xu et al., 2012, Iacaruso et al., 2017).

Location-dependent plasticity rules vary along the proximodistal axis of dendrites because bAPs are attenuated as they propagate away from the soma (Regehr et al., 1989, Spruston et al., 1995, Golding et al., 2002, Waters et al., 2003, Froemke et al., 2005, Sjostrom and Hausser, 2006). In distal dendritic compartments, smaller amplitude bAPs evoke less voltage-dependent calcium-influx, such that induction protocols that usually evoke LTP result in LTD (Nevian and Sakmann, 2006, Sjostrom and Hausser, 2006). Experimentally boosting distal bAP amplitude can restore LTP by amplifying bAP-dependent calcium influx (Sjostrom and Hausser, 2006), suggesting that location-dependent variation in synaptic plasticity rules is primarily determined by the local amplitude and timing of bAP-dependent calcium influx.

In principle, bAP-dependent calcium influx can also vary in a branch-specific manner, independent of distance from the soma. Dendritic bAP-dependent calcium influx could be shaped by local variability in ion channel expression, synaptic inputs, or electrotonic impedance due to dendritic branch structure (Rall and Rinzel, 1973, Vetter et al., 2001, Frick et al., 2004, Losonczy et al., 2008, Yaeger et al., 2019). Branch-specific variation in bAP-dependent calcium influx provides a potential mechanism for diversifying synaptic tuning across the dendritic tree, which is a core feature of computational models of neurons as hierarchical nonlinear integrators (Poirazi et al., 2003, Hausser and Mel, 2003, Francioni and Harnett, 2021, Bicknell and Hausser, 2021). The existence of branch-specific variation in bAP-dependent calcium influx would also affect the interpretation of dendritic calcium transients imagined *in vivo,* which can be evoked by local synaptic integration or global bAP-dependent influx (Wilson et al., 2016, Iacaruso et al., 2017, Yaeger et al., 2019, Beaulieu-Laroche et al., 2019, Kerlin et al., 2019, Francioni et al., 2019).

To discover potential dendrite branch-specific and distance-independent variation in bAP amplitude and bAP-dependent calcium influx, we measured bAP-dependent calcium influx in multiple dendrites of individual cortical L2/3 pyramidal cells in mice. We used somatic and dendritic electrical recordings, two-photon calcium imaging and glutamate uncaging, and dendritic voltage-imaging to experimentally identify dendrite branch-specific changes in bAP amplitude and its consequence for bAP-dependent calcium influx. We discovered branch-specific reductions in bAP amplitude that are independent of distance from the soma. In these branches, bAPs fail to evoke calcium influx through voltage-gated calcium channels (VGCCs), but still amplify synaptically-mediated calcium influx through NMDA receptors (NMDARs). We demonstrated that these branches successfully propagate bAPs in the absence of synaptic input and contain VGCCs, but exhibit more elaborate dendritic branching, which decreases their electrotonic impedance and reduces the amplitude of the bAP. Our results reveal branch-specific variation in bAP-dependent calcium signals, providing an additional mechanism by which calcium-dependent plasticity induction may vary throughout individual neurons.

## Results

We measured calcium influx in the apical dendrites of cortical L2/3 pyramidal cells to investigate whether voltage-dependent calcium signals are regulated in a branch-specific manner. We acquired whole-cell current-clamp recordings from individual cells filled through the recording pipette with 10 μM Alexa Fluor 594 and 300 μM Fluo-5F to visualize neuronal morphology and monitor changes in intracellular calcium concentration using red and green fluorescence, respectively (Carter and Sabatini, 2004). Calcium-dependent fluorescence transients were measured as the change in green relative to red fluorescence *(ΔG/R),* which is linearly proportional to Δ*Ca* and comparable across recordings (Carter and Sabatini, 2004). We found that bAP-evoked calcium influx *(ΔCa_AP_*) varies across dendritic spines from within the same cells, even when matched for distance from the soma (Figure 1). Across many neurons and recording sites, we observed that Δ*Ca_AP_* was highly correlated in nearby spines (<6 μm) and between spines and their parent dendritic shafts (Figure 1 – figure supplement 1), indicating that variations in Δ*Ca_AP_* are regulated across dendritic branches, rather than across individual dendritic spines. These data suggest that voltage-gated calcium channels (VGCCs), which mediate bAP-evoked calcium influx and are responsible for various forms of plasticity (Kapur et al., 1998, Dolmetsch et al., 2001, Yasuda et al., 2003, Nevian and Sakmann, 2006, Scheuss et al., 2006, Bloodgood and Sabatini, 2007, Brigidi et al., 2019), can be differentially activated by bAPs in a branch-specific manner within individual L2/3 pyramidal cells.

**Figure 1.**
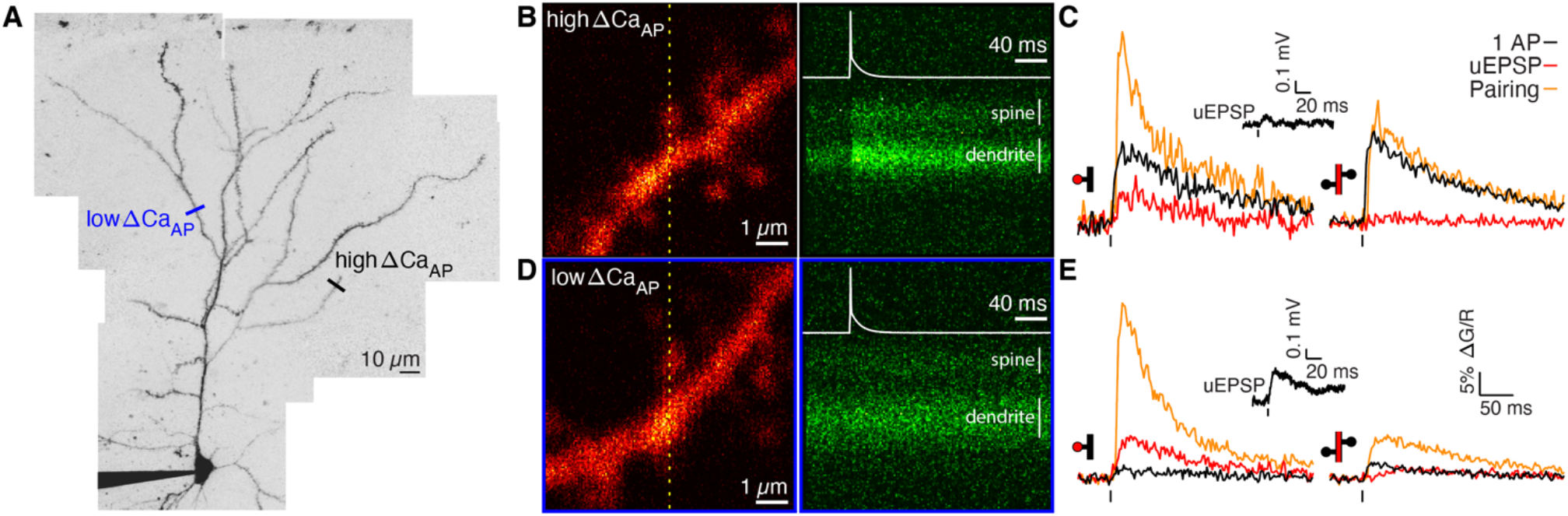
Voltage-dependent calcium influx varies in a branch-specific manner throughout L2/3 pyramidal cells. A) Maximum z-projection of Alexa 594 fluorescence showing the full apical dendritic morphology of a cortical L2/3 pyramidal cell. Two sites corresponding to the dendritic regions with low (blue) and high (black) ΔCa_AP_ are indicated. B) Frame scan (left) showing high ΔCa_AP_ branch. Dotted yellow line indicates the orientation of the line scan used to acquire the data in the kymograph (right) of Fluo-5F fluorescence in the spine and neighboring dendrite evoked by a bAP. White vertical lines indicate ROIs for spine and dendrite. Inset: bAP waveform recorded at the soma. C) Average calcium-dependent fluorescence transients in a high ΔCa_AP_ dendritic spine (left) and parent dendrite (right) evoked by 1 bAP, an uEPSP, and pairing of an uEPSP with 1 bAP (bAP evoked 5 ms after the uEPSP). Inset: average somatic whole-cell recording of the uEPSPs. In this and all presentations of imaging data, the red shaded area of the spine and dendritic schematic indicates the region from which fluorescence was measured. D) As in B for the low ΔCa_AP_ branch. E) As in C for the low ΔCa_AP_ branch. Note the difference in ΔCa_AP_ (black) between the spines in B-C and D-E.

To investigate the mechanisms of branch-specific variation in voltage-dependent calcium signals, we measured calcium influx in response to glutamate uncaging-evoked excitatory postsynaptic potentials (uEPSPs) as a proxy for synaptically-evoked calcium influx *(ΔCa_uEPSP_*). In addition, we measured bAP-dependent amplification of synaptically-evoked calcium influx due to the transient relief of the Mg^2+^ block of NMDARs using uEPSP-bAP pairings *(ΔCa_pairing_*). In all uEPSP-bAP pairings, we delayed the bAP by 5 ms relative to the glutamate uncaging laser pulse to maximize bAP-dependent amplification of postsynaptic calcium influx and mimic protocols that robustly induce NMDAR-dependent LTP (Froemke et al., 2005, Nevian and Sakmann, 2006, Holbro et al., 2010). Spines on branches with and without bAP-evoked calcium influx showed Δ*Ca_uEPSP_* and supralinear bAP-dependent amplification in Δ*Ca_pairing_* (Figure 1B-E) (Holbro et al., 2010), even though one of the spines exhibited almost no bAP-evoked calcium influx when the bAP was evoked by itself (Figure 1D-E). These data suggest that bAP-evoked calcium influx through VGCCs and bAP-dependent amplification of synaptically-evoked calcium influx through NMDARs can be regulated independently in a branch-specific manner (Bloodgood and Sabatini, 2007, Holbro et al., 2010).

### bAP-evoked calcium influx and bAP-dependent amplification are decoupled in L2/3 pyramidal cells

To systematically investigate the regulation of bAP-evoked calcium influx and bAP-dependent amplification of synaptically-evoked calcium influx, we recorded Δ*Ca_AP_*, Δ*Ca_uEPSP_*, and Δ*Ca_pairing_* in 97 dendritic spines from 33 cells spanning the full proximodistal extent of the dendrites. To focus our analysis on the supralinear component of Δ*Ca_pairing_*, which is primarily carried by NMDARs (Holbro et al., 2010), we measured amplification as Δ*Ca_amp_* = Δ*Ca_pairing_ — (ΔCa_AP_* + Δ*Ca_uEPSP_*) (Figure 2A). We found that Δ*Ca_AP_* and Δ*Ca_amp_* were uncorrelated across dendritic spines in L2/3 pyramidal cells (Figure 2B). Based on the amplitude of bAP-evoked calcium influx, we grouped the data into high Δ*Ca_AP_* (0.1 < Δ*Ca_AP_* < 0.3, selected to avoid outliers), and low Δ*Ca_AP_ (ΔCa_AP_* < 0.04) populations (Figure 2B) and plotted the average fluorescence transients in each to highlight the divergence between Δ*Ca_AP_* and Δ*Ca_amp_* (Figure 2C-D). This selection criterion is used throughout all figures except for Figure 5B-D. Low Δ*Ca_AP_* branches exhibited almost no calcium influx evoked by bAPs but still exhibited large bAP-dependent amplification (Figure 2C-D). Δ*Ca_AP_* and Δ*Ca_amp_* showed similar patterns in dendritic spines and parent dendritic shafts (Figure 2C-D, insets). We found that although Δ*Ca_AP_* attenuates with distance from the soma (Figure 2E), it exhibits significant variability in distal branches, such that pairs of dendritic spines that are on different branches within the same neuron but at a similar distance from the soma sometimes had large differences in calcium influx (Figure 2E-F). Δ*Ca_amp_* varied over a similar range at all distances from the soma (Figure 2 – figure supplement 1A). The divergence between Δ*Ca_AP_* and Δ*Ca_amp_* was not related to the degree of NMDAR activation (Figure 2 – figure supplement 1B) nor the uEPSP amplitude or time-course (Figure 2 – figure supplement 1C-E). Furthermore, we observed low Δ*Ca_AP_* branches in the presence of 10 μM SR 95531, an antagonist of ionotropic GABA receptors, indicating that low Δ*Ca_AP_* branches are not caused by inhibition (data not shown). Together, these data indicate that Δ*Ca_AP_* and Δ*Ca_amp_* are regulated independently in the dendrites of L2/3 pyramidal cells.

**Figure 2.**
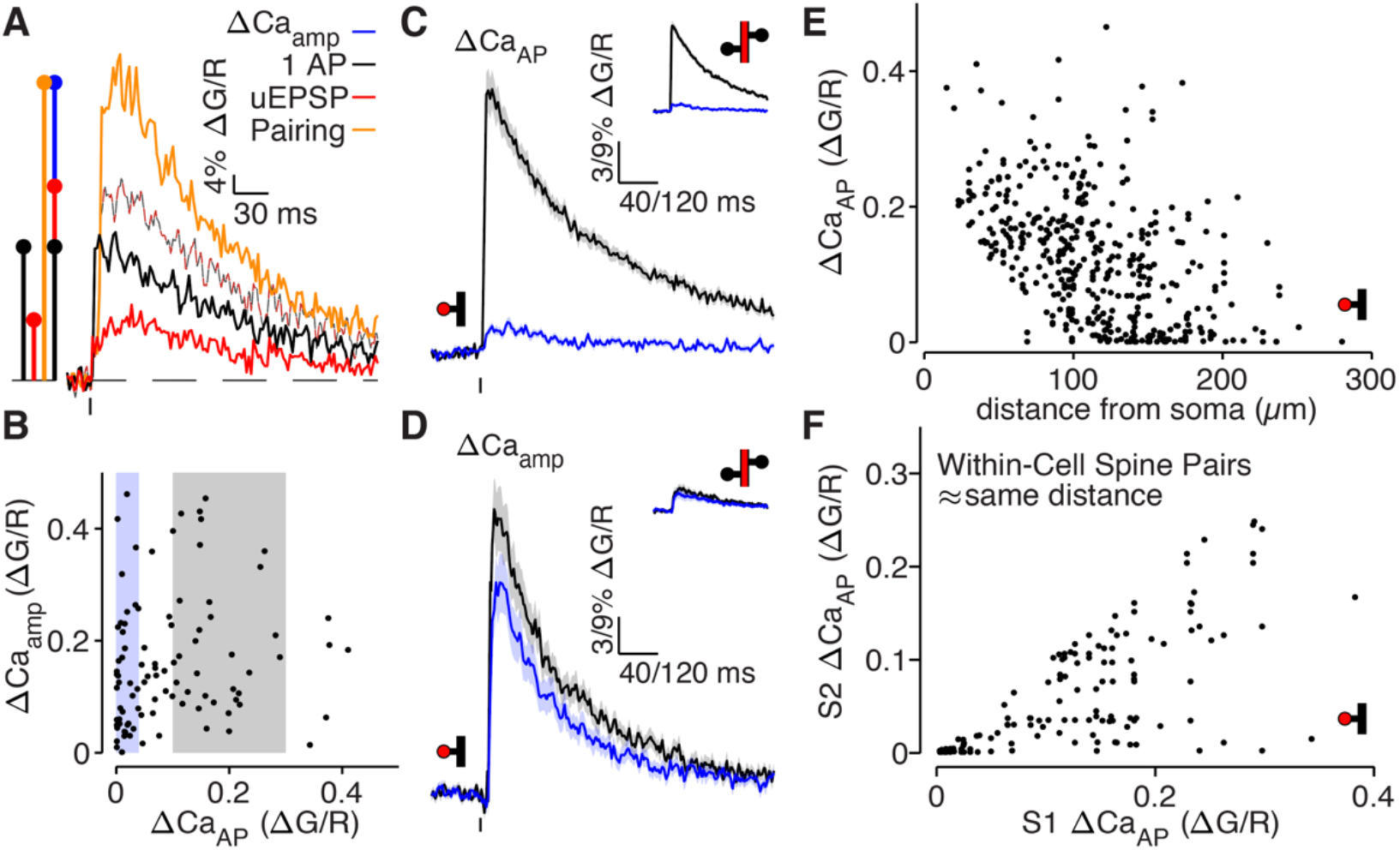
bAP-evoked calcium influx and bAP-dependent amplification are decoupled in L2/3 pyramidal cells. A) Average calcium-dependent fluorescence transients recorded from a dendritic spine and a schematic of the method for computing the amplification of calcium influx caused by interaction of the bAP and uEPSP, which results in additional influx (blue) relative to the sum of the bAP and uEPSP components measured independently (black/red). B) bAP-dependent amplification vs. bAP-evoked calcium influx. Shaded areas indicate selection of high ΔCa_AP_ (gray) and low ΔCa_AP_ (blue) spines used in panels C and D. C) ΔCa_AP_ in high (black) and low (blue) ΔCa_AP_ dendritic spines and parent dendritic shafts (inset). In this and all presentations of average fluorescence transients, the line and shaded areas show the mean and standard error around the mean. D) ΔCa_amp_ in high (black) and low (blue) ΔCa_AP_ dendritic spines and parent dendritic shafts (inset). E) ΔCa_AP_ across distance from soma in all dendritic spines. F) Comparison of ΔCa_AP_ between pairs of dendritic spines (i.e. S1 & S2) that were recorded from the same cell, on different dendritic branches, and at a similar distance from the soma (<20 μm relative distance from soma). Each pair was sorted so the abscissa indicates the ΔCa_AP_ of the spine with higher calcium influx.

We considered three distinct mechanistic explanations for why low Δ*Ca_AP_* branches exhibit bAP-dependent amplification (Δ*Ca_amp_*) but fail to exhibit bAP-evoked calcium influx (Δ*Ca_AP_*):

1. Low Δ*Ca_AP_* branches contain fewer (or no) voltage-gated calcium channels (VGCCs).
2. bAPs fail to backpropagate into low Δ*Ca_AP_* branches without the support of EPSPs.
3. bAP amplitude in low Δ*Ca_AP_* branches does not exceed the threshold for VGCC opening.

### Low Δ*Ca_AP_* branches contain voltage-gated calcium channels

Low Δ*Ca_AP_* branches may contain functional VGCCs that are not opened by a single bAP but do open in response to stronger depolarizations. To examine this possibility, we imaged calcium influx evoked by a burst of 5 bAPs at 150 Hz in dendritic shafts located >75 μm from the soma L2/3 pyramidal cells (Figure 3A) (Larkum et al., 2007). In most low Δ*Ca_AP_* branches, calcium influx could be evoked by a burst of 5 bAPs (Figure 3A-B). However, the calcium influx evoked by a burst of 5 bAPs in high Δ*Ca_AP_* branches was higher than in low Δ*Ca_AP_* branches (Figure 3A-B). These data are consistent with the presence of VGCCs in low Δ*Ca_AP_* branches, although they do not distinguish whether there is a lower density of VGCCs (or a higher voltage threshold), or if bursts of bAPs are less effective at depolarizing low Δ*Ca_AP_* branches.

**Figure 3.**
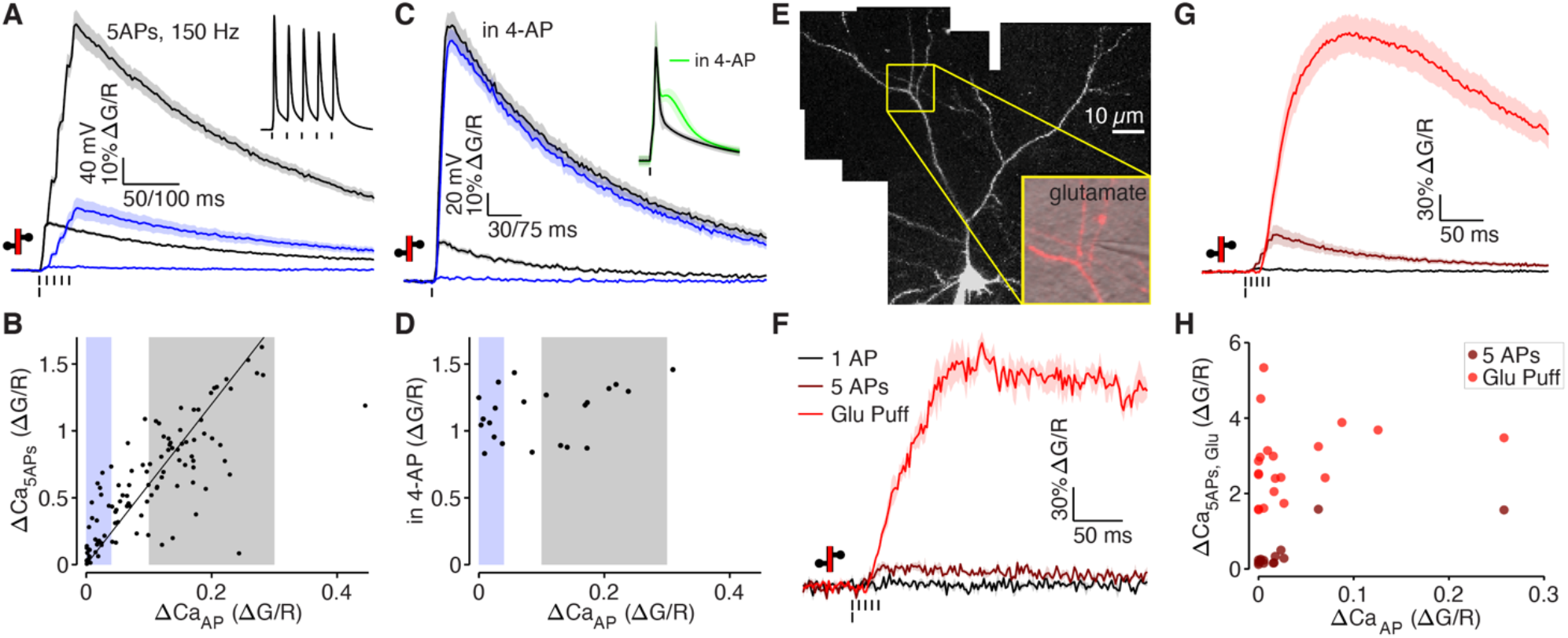
Low ΔCa_AP_ dendrites contain voltage-gated calcium channels. A) Calcium-dependent fluorescence transients evoked by 1 bAP or 5 bAPs at 150 Hz in high (black) and low (blue) ΔCa_AP_ dendrites. Inset: example trace of somatic whole-cell recording during 5 bAP burst protocol. B) Comparison of peak calcium influx evoked by 1 bAP or 5 bAPs in each dendrite. Gray and blue patches indicate selection of high (gray) and low (blue) ΔCa_AP_ spines used in panel A. The line y = 6x is plotted for reference. C) Calcium-dependent fluorescence transients evoked by 1 bAP before and after application of 2 mM 4-AP in high (black) and low (blue) ΔCa_AP_ branches. Inset: average AP waveform before and after application of 4-AP. D) Comparison of peak calcium influx evoked by 1 bAP before and after application of 2 mM 4-AP. Gray and blue patches indicate selection of high (gray) and low (blue) ΔCa_AP_ spines used in panel C. E) Maximum z-projection of Alexa 594 fluorescence from a L2/3 pyramidal cell with differential-interference contrast image overlaid showing glutamate puffing pipette. The full apical dendritic morphology was not imaged. F) Calcium-dependent fluorescence transients evoked by 1 bAP, 5 bAPs at 150 Hz, and a 1 mM glutamate puff in the example branch from panel E. G) Average calcium-dependent fluorescence transients evoked by 1 bAP, 5 bAPs at 150 Hz, and a 1 mM glutamate puff across all branches recorded. H) Comparison of peak calcium influx evoked by bAP, 5 bAPs at 150 Hz, and a 1 mM glutamate puff in all branches. The response to 5 bAPs was not recorded in all dendrites.

We attempted to make the depolarization evoked across the dendritic tree more uniform by triggering bAPs in the presence of the potassium channel antagonist 4-AP (Figure 3C-D) (Hoffman et al., 1997). In all somatic recordings, application of 2 mM 4-AP increased the after-depolarization of the bAP (Figure 3C, inset), confirming the efficacy of the drug. Applying 4-AP significantly increased Δ*Ca_AP_* in every branch, regardless of how much calcium influx was evoked in control conditions (Figure 3C-D). Because Δ*Ca_AP_* evoked in the presence of 4-AP is similar in high and low Δ*Ca_AP_* branches (Figure 3C), we conclude that VGCCs are present in all dendritic compartments and mediate substantial calcium influx in response to sufficiently large depolarizations.

To determine if VGCCs can be effectively opened by local depolarization in low Δ*Ca_AP_* branches, we measured calcium influx evoked by direct application of 1 mM glutamate delivered via a patch pipette using a picospritzer (Figure 3E-H). We added 20 μM CPP and 50 μM MK-801 to the puff and bath solutions to block NMDARs and limit our measurements to calcium influx mediated by VGCCs. In every dendrite examined, direct application of glutamate led to significant calcium influx, regardless of Δ*Ca_AP_* (Figure 3F-H). The calcium signal was not an artifact of mechanical displacement due to the picospritzer (Figure 3 – figure supplement 1A-B) and did not include an NMDAR component (Figure 3 – figure supplement 1C-D), consistent with calcium influx mediated by VGCCs. We note that the large calcium signals evoked by glutamate demonstrate that the similarity in bAP-evoked calcium influx between high and low Δ*Ca_AP_* measured in 4-AP was not due to saturation of the calcium indicator. Collectively, our data indicate that all dendritic branches contain VGCCs, which can be opened by local depolarization. However, a specific population of low Δ*Ca_AP_* branches exhibit little to no calcium influx in response to back-propagating bAPs.

### bAPs propagate to low Δ*Ca_AP_* branches in the absence of EPSPs

We only observe significant bAP-dependent calcium influx in low Δ*Ca_AP_* branches if the bAP is paired with synaptic input evoked by glutamate uncaging. Despite the small size of the uEPSP, it is possible that bAPs only back-propagate to low Δ*Ca_AP_* branches when assisted by local depolarization (Magee and Johnston, 1997, Gasparini et al., 2007). Since most of the depolarization evoked by EPSPs is carried by AMPARs, we measured Δ*Ca_AP_*, Δ*Ca_uEPSP_*, and Δ*Ca_pairing_* after applying the selective AMPAR antagonist, NBQX (Figure 4A-B). We observed bAP-dependent amplification of the remaining NMDAR-mediated calcium influx in all dendritic spines examined, despite fully blocking the depolarization evoked by glutamate uncaging (Figure 4A-B). As expected, glutamate uncaging evokes some calcium influx via NMDARs in the presence of NBQX, since NMDARs are only partially blocked by Mg^2+^ at resting potentials (Jahr and Stevens, 1990) and the driving force on calcium is extremely high. These data suggest that bAPs propagate to low Δ*Ca_AP_* branches without an uEPSP-mediated local depolarization.

**Figure 4.**
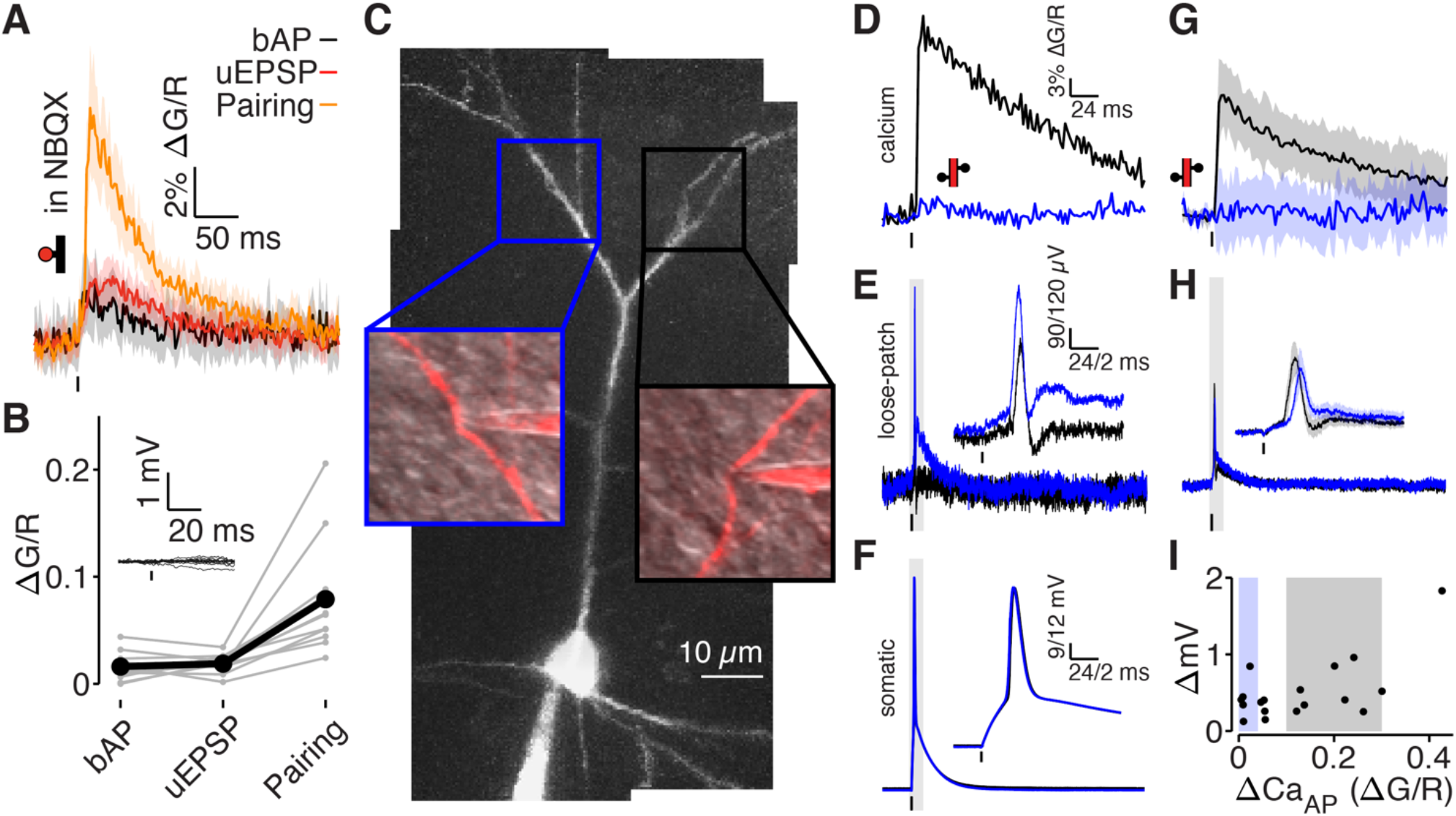
bAPs propagate to low ΔCa_AP_ dendrites in the absence of EPSPs. A) Calcium-dependent fluorescence transients evoked by bAPs, uEPSPs, and bAP/uEPSP pairings in the presence of 10 μM NBQX. B) Comparison of peak calcium influx in spines evoked by bAP, uEPSP, and bAP/uEPSP pairing, in the presence of 10 μM NBQX. Inset: whole-cell recordings from glutamate uncaging in the presence of NBQX. C) Maximum z-projection of Alexa 594 fluorescence from a L2/3 pyramidal cell with a whole-cell recording in the soma and consecutive loose-patch recordings in two branches. Insets: red fluorescence overlaid on differential interference contrast image of loose-patch configuration for each dendrite. The full apical dendritic morphology was not imaged. D) Calcium-dependent fluorescence transients evoked by bAPs in each dendrite from panel C, recorded during loose-patch recordings. E) Electrical signals evoked by bAPs measured with dendritic loose-patch recordings in each dendrite from panel C. Inset: expanded trace from gray shaded region. F) Somatic whole-cell recording of bAPs during each loose-patch recording from panel C. Inset: expanded trace from gray shaded region. G) Average calcium-dependent fluorescence transients evoked by bAPs in high (black) and low (blue) ΔCa_AP_ branches during loose-patch recordings. H) Average electrical signal evoked by bAPs in high (black) and low (blue) ΔCa_AP_ branches measured with loose-patch recordings. I) Comparison of peak calcium influx and peak electrical signal evoked in each dendrite by bAPs. Patches indicate selection of high (gray) and low (blue) ΔCa_AP_ branches used in panels G and H.

To test whether bAPs propagate to low Δ*Ca_AP_* branches in the absence of glutamate receptor activation, we acquired loose-patch recordings in current-clamp from 19 dendrites in 15 neurons while simultaneously recording from the soma to measure the dendritic electrical signal evoked by a bAP (Figure 4C-I). Simultaneous two-photon imaging was used to monitor dendritic calcium influx. All recording sites were ≥75 μm from the soma. In the neuron shown in Figure 4C, we acquired consecutive loose-patch recordings from one high Δ*Ca_AP_* branch and one low Δ*Ca_AP_* branch. Only the high Δ*Ca_AP_* branch had detectable Δ*Ca_AP_,* but both dendritic recordings exhibited clear electrical signals evoked by the bAP (Figure 4D-F). We observed bAP-evoked electrical signals in all dendrites, even though only some exhibited Δ*Ca_AP_* (Figure 4G-I). These loose-patch recordings do not permit direct measurement of intracellular bAP amplitude (Figure 4 – figure supplement 1); however, they demonstrate that bAPs propagate to low Δ*Ca_AP_* branches without the support of EPSPs.

### bAP amplitude is attenuated in low Δ*Ca_AP_* branches

To measure the intracellular dendritic voltage evoked by bAPs, we performed interleaved dendritic voltage and calcium imaging while performing somatic whole-cell current-clamp recordings (Figure 5). Voltage imaging was achieved by expressing a highly-sensitive near-infrared genetically-encoded voltage indicator, QuasAr6a (Tian et al., 2021). Both voltage and calcium signals were recorded via structured illumination one-photon microscopy. All recording sites were ≥75 μm from the soma (except for the closest site in panel A). Consistent with our results from Figure 4, we observed bAP-dependent voltage signals in all branches, including those that did not exhibit bAP-evoked calcium influx (Figure 5). To compare high and low Δ*Ca_AP_* branches, we adjusted our selection criterion to account for differences in the range of Δ*Ca_AP_* measured with two-photon and one-photon calcium imaging (the selection range described above was scaled by 1.75, Figure 5B-D). Although low Δ*Ca_AP_* branches exhibited clear bAP-evoked voltage waveforms (Figure 5C), the amplitudes of the dendritic bAP-evoked voltage waveforms were attenuated in low Δ*Ca_AP_* branches (Figure 5C-D). These data demonstrate that bAP amplitude is reduced in low Δ*Ca_AP_* branches.

**Figure 5.**
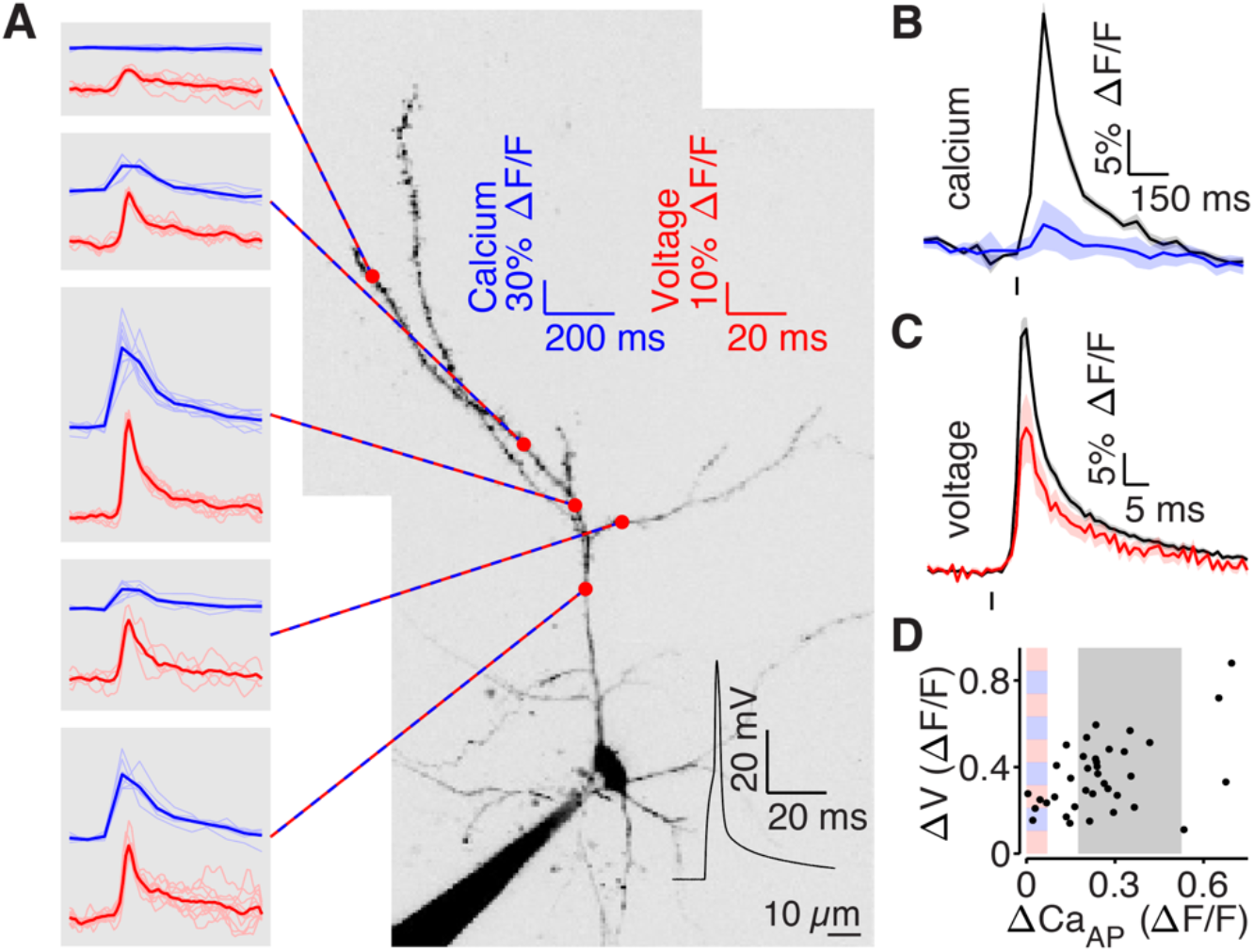
bAP amplitude is attenuated in low Δ*Ca_AP_* branches. A) Mean calcium-dependent (blue) and voltage-dependent (red) fluorescence transients evoked by 1 bAP (left) measured in the 5 branches indicated in the maximum z-projection of Alexa 594 fluorescence from a L2/3 pyramidal cell (right). The black trace shows the bAP waveform recorded from the soma during calcium and voltage imaging. Note that calcium- and voltage-dependent transients are shown on ~10-fold different time scales. The full apical dendritic morphology was not imaged. B) Average calcium-dependent fluorescence transient in high (black) and low (blue) ΔCa_AP_ branches evoked by 1 bAP. C) Average voltage-dependent fluorescence transient in high (black) and low (red) ΔCa_AP_ branches evoked by 1 bAP. D) Comparison of the peak voltage- and calcium-dependent fluorescence transient evoked by 1 bAP. Shaded regions indicate selection criterion for high (gray) and low (blue/red) ΔCa_AP_ branches.

Collectively, our data support the conclusion that low Δ*Ca_AP_* branches exhibit less bAP-evoked calcium influx because of branch-specific reductions in local bAP amplitude. First, our experiments with bursts of 5 bAPs, application of 4-AP, and glutamate application show that low Δ*Ca_AP_* branches contain VGCCs that open in response to large depolarizations (Figure 3). Second, our experiments with NBQX and dendritic loose-patch recordings show that bAPs successfully propagate to low Δ*Ca_AP_* branches in the absence of synaptic input (Figure 4). Finally, our voltage imaging experiments show that the amplitude of bAPs is attenuated in low Δ*Ca_AP_* branches (Figure 5). These data are consistent with the mechanistic explanation that bAPs fail to evoke calcium influx in low Δ*Ca_AP_* branches because the local bAP amplitude does not exceed the threshold for VGCC opening. Therefore, we investigated why bAPs are selectively attenuated in low Δ*Ca_AP_* branches.

### bAP-evoked calcium influx attenuates at branch points

bAP amplitude and bAP-evoked calcium influx can attenuate at branch points due to dendritic impedance mismatch and localized expression of potassium channels (Spruston et al., 1995, Williams and Stuart, 2000, Vetter et al., 2001, Frick et al., 2003, Cai et al., 2004, Gasparini et al., 2007, Harnett et al., 2013). Consistent with these observations, we found that Δ*Ca_AP_* often attenuates across branch points, as revealed by comparing differences in Δ*Ca_AP_* at pairs of sites within the same dendritic segment or across one dendritic branch point (Figure 6 – figure supplement 1A-C). The ratio of Δ*Ca_AP_* (influx at distal site divided by proximal site) tends to be smaller for pairs of sites spanning a branch point than for those within the same segment (rank-sum: *p=0.0145,* Figure 6 – figure supplement 1C). This result was corroborated by the observation that low Δ*Ca_AP_* branches typically had higher dendritic branch order (defined as the number of on-path branch points between the soma and the recording site) than high Δ*Ca_AP_* branches (Figure 6 – figure supplement 1D). While these data suggest a relationship between Δ*Ca_AP_* and the pattern of dendritic branching, they do not fully account for the variance in Δ*Ca_AP_* observed across branches in L2/3 pyramidal cells (Figure 6 – figure supplement 1D, R^2^=0.178).

### Low Δ*Ca_AP_* branches have a more elaborate branch structure than high Δ*Ca_AP_* branches

The electrotonic impedance (i.e., passive input resistance) of a dendrite scales inversely with the surface area of membrane. Therefore, if low Δ*Ca_AP_* branches are within sections of the dendritic tree that contain more extensive branching, then they have a smaller impedance, which might reduce bAP amplitude and Δ*Ca_AP_*. To examine this possibility, we developed a metric for impedance based on the dendritic morphology (“branch impedance”) that accounts for all nearby dendritic branches and is independent of branch order. This approach measures the total length of dendrite near a recording site with an exponential filter to discount branches that are further from the site. We measured branch impedance over a length constant of λ=145 μm because propagating bAPs simultaneously depolarize ~145 μm of dendrite in L2/3 pyramidal cells (conduction velocity is 154.9 μm/ms and AP duration is 0.94 ms, Figure 6 – figure supplement 2).

We compared the branch impedance of pairs of dendritic recording sites within the same cell that were located approximately the same distance from the soma but for which there was a large difference in Δ*Ca_AP_* (Figure 6A-E). The low Δ*Ca_AP_* site had a smaller branch impedance than the high Δ*Ca_AP_* site in all pairs (Figure 6E), indicating that low Δ*Ca_AP_* sites are surrounded by more extensive branching than high Δ*Ca_AP_* sites. To confirm that branch impedance scales with input resistance, we reconstructed all cells used in this analysis as biophysical compartment models in the NEURON simulation environment and directly measured the input resistance by injecting small hyperpolarizing currents into each recording site (Carnevale and Hines, 2006). In every pair, the low Δ*Ca_AP_* site had a smaller input resistance than the high Δ*Ca_AP_* site (Figure 6F). Additionally, branch impedance was highly correlated with input resistance (R^2^ = 0.93 for λ=145 μm), confirming its validity as a measure of electrotonic properties (Figure 6 – figure supplement 3A-B). These results and conclusions hold for measurements of branch impedance over length constants ranging from 5 – 400 μm (Figure 6 – figure supplement 3C).

**Figure 6.**
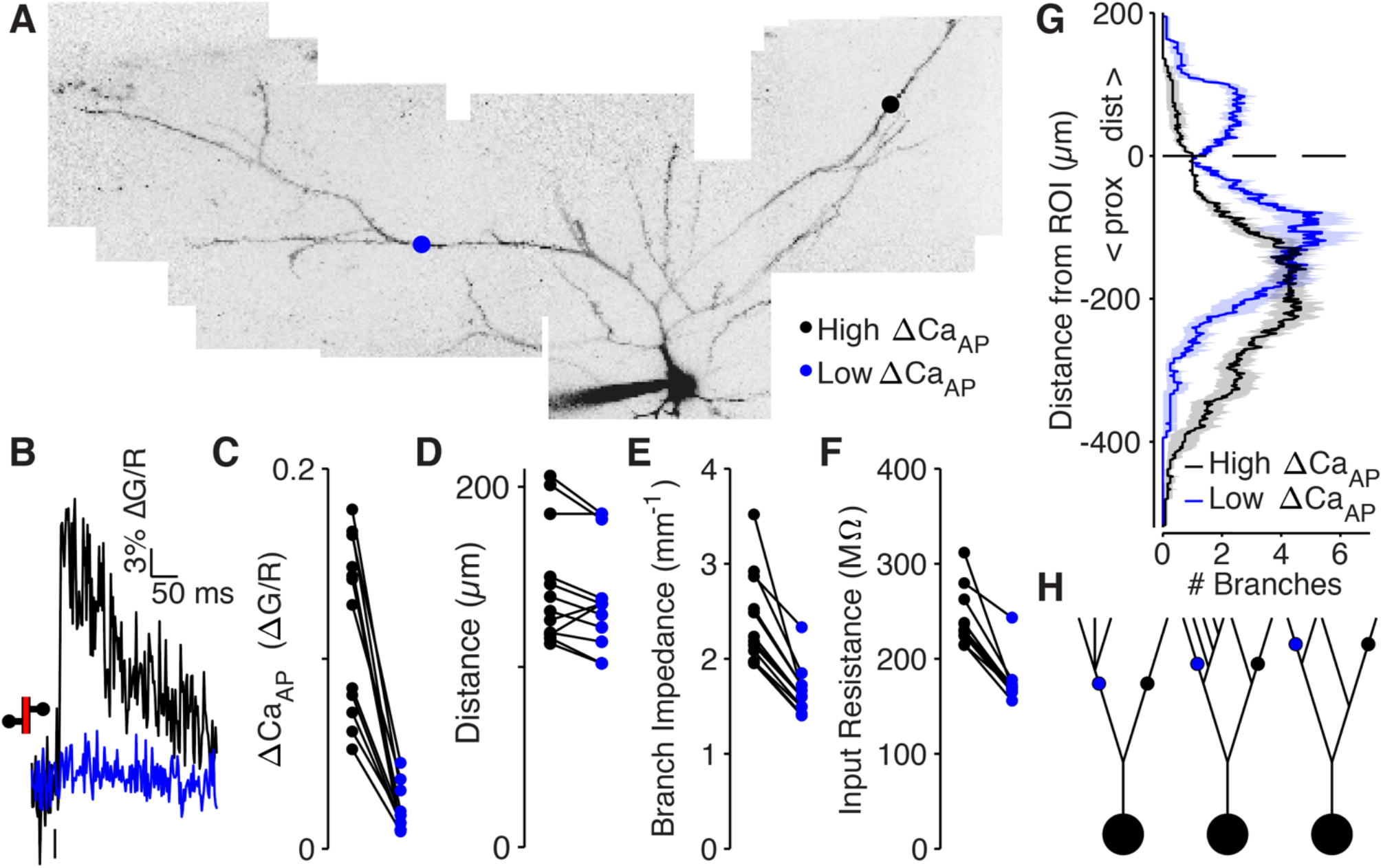
Low ΔCa_AP_ dendrites have a more elaborate branch structure than high ΔCa_AP_ dendrites. A) Maximum z-projection of Alexa 594 fluorescence from a L2/3 pyramidal cell. Dots indicate location of calcium imaging sites for a high (black) and low (blue) ΔCa_AP_ branch. The two sites are approximately the same distance from the soma (high ΔCa_AP_: 145μm, low ΔCa_AP_: 131μm). B) Calcium-dependent fluorescence transient evoked by bAP in high (black) and low (blue) ΔCa_AP_ sites from panel A. C) Comparison of ΔCa_AP_ in distance-matched, within-cell pairs. Ratio: 0.17, (range: 0.06-0.29). D) Comparison of the distances from the soma for each pair of recording sites. E) Comparison of branch impedance in high (black) and low (blue) ΔCa_AP_ sites, using a distance-discounted measurement of nearby dendritic branches. F) Comparison of input resistance for high (black) and low (blue) ΔCa_AP_ sites shown in panels C and D, measured in computational simulations of compartment models of each cell in NEURON. G) Average number of branches at a given distance from a recording site for high (black) and low (blue) ΔCa_AP_ dendrites. This curve was multiplied by a symmetric exponential filter with a length constant of 145 μm and then integrated to compute the branch impedance in panel E. Mean ± SEM. H) Schematic of morphologies with same branch order but different branch impedance. The blue and black site in each neuron have the same branch order, but the branch impedance of the blue site is lower due to the number of branches distal to the site (left), the number of branches on a sister dendrite (middle), and the distance from a previous branch (right).

To inspect why low Δ*Ca_AP_* branches had a smaller branch impedance, we plotted the average number of dendritic branches distal and proximal to the target site (Figure 6G). Low Δ*Ca_AP_* branches are immediately proximal to dendritic branch elaboration (e.g., Figure 6A, c.f. with Figure 6H, left), and closer to sister dendrites emerging proximal to the measurement site (c.f. with Figure 6H, middle and right). These results demonstrate that low Δ*Ca_AP_* branches are surrounded by more elaborate branching patterns than high Δ*Ca_AP_* branches, suggesting that dendritic branch structure may be sufficient to selectively reduce the amplitude of bAPs.

### Dendritic branch structure is sufficient to explain reductions in bAP-evoked calcium influx

To determine if dendritic branch structure is sufficient for reducing Δ*Ca_AP_*, we reproduced our experimental results using a biophysical compartment model. We held all physiological parameters constant throughout each model except for the dendritic branch structure, allowing us to specifically measure the effects of dendritic morphology between high and low Δ*Ca_AP_* branches (Vetter et al., 2001). Using NEURON compartmental models of 8 reconstructed L2/3 pyramidal cells (all from the pairwise analysis described in Figure 6), we evoked a bAP with somatic current injection and measured the intracellular voltage waveform and the resulting calcium conductance at the same sites in which we had imaged calcium influx in acute brain slices (Figure 7A-E). For each cell, we fit the density of dendritic A-Type potassium channels such that the bAP peaked at −10 mV in the high Δ*Ca_AP_* site (Figure 7D), which is consistent with the amplitude of bAPs measured ~100 μm from the soma in layer 2-3 pyramidal cells with dendritic whole-cell recordings (Larkum et al., 2007, Smith et al., 2013). Under these conditions, for every within-cell pair, the low Δ*Ca_AP_* site had smaller bAP amplitude and peak calcium conductance, despite having the same channel densities as the high Δ*Ca_AP_* site (Figure 7B-E). As an additional measure of dendritic excitability, we plotted the A-Type potassium channel density required for bAPs to peak at −10 mV in each dendritic site (Figure 7F). Low Δ*Ca_AP_* sites required lower potassium channel densities for their bAPs to peak at −10 mV, indicating that the branch structure surrounding low Δ*Ca_AP_* sites is less excitable than for high Δ*Ca_AP_* sites (Figure 7F). These data demonstrate that dendritic branch structure is sufficient to impact Δ*Ca_AP_* by altering the dendritic bAP amplitude in a branch-specific manner.

**Figure 7.**
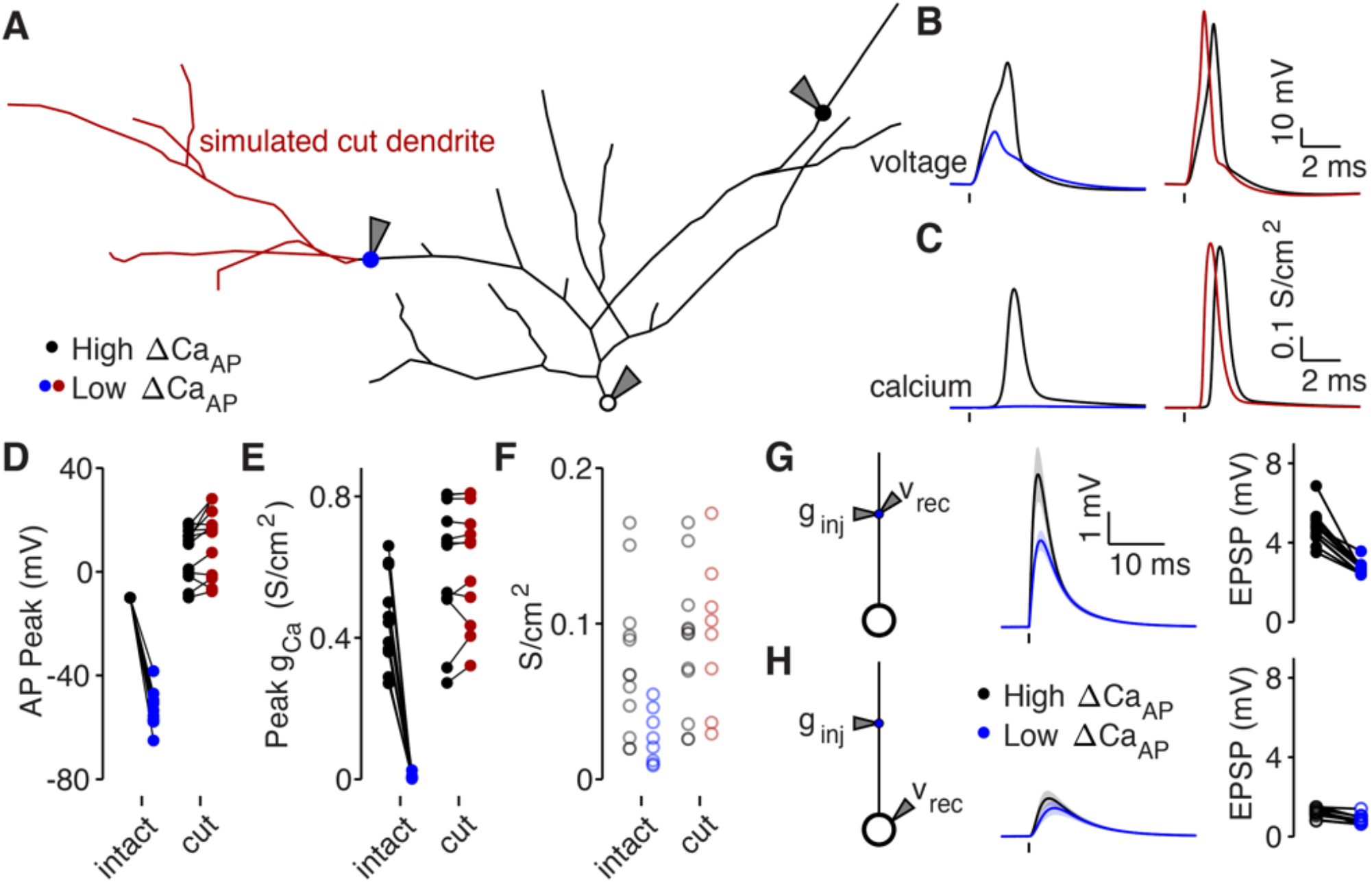
Dendritic branch structure is sufficient to explain reductions in bAP-evoked calcium influx. A) Schematic of a neuron morphology used for NEURON simulations derived from the neuron shown in Figure 6A. Blue and black dots indicate high (black) and low (blue) ΔCa_AP_ recording sites. Red dendrites indicate section of dendrite that was computationally “cut” in a subset of experiments. Recordings and injections were performed at the sites indicated with the gray triangles. B) Dendritic voltage recordings of bAPs from high (black) and low (blue/red) ΔCa_AP_ sites in panel A in NEURON for an intact cell (left) or with cut dendrites (right). C) Calcium conductances recorded from high (black) and low (blue/red) ΔCa_AP_ sites in panel A evoked by a back-propagating bAP in NEURON for an intact cell (left) or with cut dendrites (right). D) Comparison of peak bAP voltage for each high/low ΔCa_AP_ pair recorded in NEURON from same sites as experimental recordings, for intact and cut cells. Dendritic potassium channel density was fit so the high ΔCa_AP_ site would have a peak of −10mV in intact cells. Rank-sum – Intact: p=0.00004, Cut: p=0.17. E) Comparison of peak calcium conductance during bAP for each high/low ΔCa_AP_ site pair, as in panel D. Rank-sum – Intact: p=0.00004, Cut: p=0.71. F) A-Type potassium channel density required for the bAP to peak at −10 mV for each site in intact or cut cells. Ranksum – Intact: p=0.009, Cut: p=0.67. G) Left: a synaptic conductance was injected into each site and the local EPSP was recorded in NEURON. Middle: dendritic EPSPs recorded in high and low ΔCa_AP_ dendritic sites. Right: comparison of dendritic EPSP amplitudes recorded in each high/low ΔCa_AP_ pair. Mean ± SEM. H) Same as in G, but with synaptic conductance injected into dendrite and recorded in the soma.

We performed a simulated “cutting” experiment in which we removed dendritic branches from the reconstruction that were closer to the low Δ*Ca_AP_* site than the high Δ*Ca_AP_* site (Figure 7A), to determine if simplifying the dendritic branch structure can recover Δ*Ca_AP_*. After cutting dendritic branches near the low Δ*Ca_AP_* site, differences in the bAP amplitude, calcium channel conductance, and the A-Type potassium channel density required to generate bAPs that peak at −10 mV were all nullified (Figure 7B-F). These data confirm that dendritic branch structure directly affects local dendritic excitability. Furthermore, these results support the model that increased dendritic branching around low Δ*Ca_AP_* recording sites is sufficient to reduce bAP amplitude and Δ*Ca_AP_*.

Branch structure-dependent changes in dendritic excitability may also affect the amplitude, kinetics, and efficacy of synaptic input arriving at each branch. Using our compartment models, we simulated synaptic input by injecting a conductance into each dendritic site (Figure 7G-H). As expected, the local EPSP amplitude was smaller in low Δ*Ca_AP_* branches because of their reduced input resistance (Figure 7G, Figure 6F). However, differences in EPSP size measured at the soma (but evoked in the dendrites) were minimal due to frequency-dependent cable attenuation (Figure 7H). The differences in local EPSP amplitude were abolished after cutting dendritic branches near the low Δ*Ca_AP_* site (Figure 7 – figure supplement 1). Therefore, although dendritic branch structure affects the local amplitude of synaptic potentials, it has a minimal impact on the efficacy of synaptic input at depolarizing the soma.

### Reductions in bAP amplitude have a selective effect on VGCC-mediated calcium influx

Our data support a model in which elaborated dendritic branching reduces dendritic excitability, which lowers bAP amplitude and bAP-evoked calcium influx. However, it remains to be determined why bAP-dependent amplification of synaptic NMDAR-mediated calcium influx is spared by reductions in bAP amplitude that impact bAP-evoked calcium influx through VGCCs. Using published data (Jahr and Stevens, 1990, Reuveni et al., 1993), we plotted the voltage-dependent open-probabilities and time constants for VGCCs and NMDARs (Figure 8A-B). Using these parameters, we simulated VGCC and NMDAR conductances and the resulting voltage-dependent calcium influx in response to quadratic depolarizing voltage stimuli designed to resemble bAPs (Figure 8C). Due to the steep voltage-sensitivity and long time constant of VGCCs (Figure 8A-B), they exhibit a dramatic dropoff of calcium influx for voltage steps peaking below −10 mV (Figure 8C-E). Although the open-probability of NMDARs is highly sensitive to peak voltage (Figure 8C-D), the interaction between open-probability and driving force minimizes differences in calcium influx evoked across a wide range of voltages (Figure 8E). These data demonstrate how small reductions in bAP amplitude can eliminate Δ*Ca_AP_* mediated by VGCCs, while largely sparing bAP-dependent amplification of synaptically-evoked, NMDAR-mediated calcium influx.

**Figure 8.**
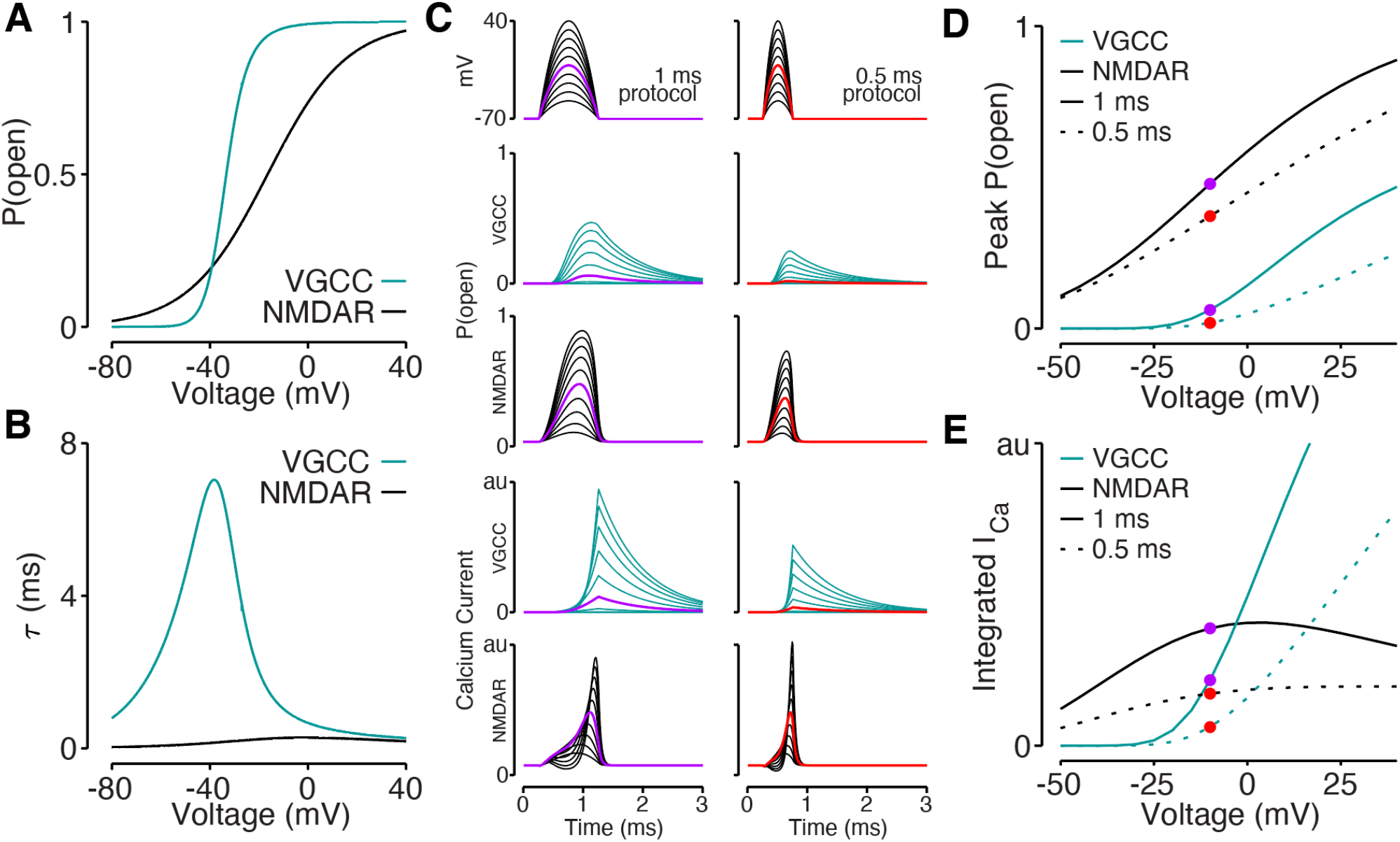
Reductions in bAP amplitude have a selective effect on VGCC-mediated calcium influx. A) Steady-state open probability of VGCCs and NMDARs as a function of voltage due to voltage-dependent activation of VGCCs and Mg^2+^-block of NMDARs. B) Voltage-dependent time constant of the activation gate for VGCCs and Mg^2+^ block for NMDARs. C) We simulated the response of VGCCs and NMDARs to a 1 ms (left) or 0.5 ms (right) quadratic depolarizing voltage with varying amplitude (1^st^ row) designed to resemble bAPs. Voltage-dependent open-probability (2^nd^ and 3^rd^ rows) and voltage-dependent calcium current (4^th^ and 5^th^ rows) for VGCCs (teal) and NMDARs (black). Purple and red traces correspond to summary data in panel D and E. D) Peak open probability of VGCCs and NMDARs during quadratic voltage protocols in panel C. E) Integral of calcium influx through VGCCs and NMDARs during quadratic voltage protocols in panel C.

## Discussion

Back-propagating APs regulate synaptic plasticity by evoking voltage-dependent calcium influx throughout dendrites. Here, we show that bAP-dependent calcium influx varies in a dendrite branch-specific manner in cortical L2/3 pyramidal cells due to branch-specific reductions in input resistance inherited from the local morphology of the dendritic tree. Sections of the dendritic tree with more elaborate branch patterns have lower input resistance, leading to a branch-specific reduction in bAP amplitude. These branches have a selective reduction in bAP-evoked calcium influx through VGCCs despite containing VGCCs and successfully propagating bAPs in the absence of synaptic input. However, these branches maintain bAP-dependent amplification of synaptically-evoked calcium influx through NMDARs due to the shallower voltage-dependence and faster kinetics of the Mg^2+^-block of NMDARs. Our results demonstrate that dendritic excitability, bAPs, and bAP-dependent calcium signals vary between branches, which may provide a mechanism for synaptic plasticity rules and the computational properties of dendrites to vary across subcompartments of individual neurons.

### Branch-specific variation in dendritic morphology shapes voltage-dependent calcium signals

Many studies have demonstrated that bAP amplitude and bAP-evoked calcium influx are attenuated in the distal compartments of pyramidal cells (Regehr et al., 1989, Spruston et al., 1995, Magee and Johnston, 1997, Waters et al., 2003, Froemke et al., 2005, Sjostrom and Hausser, 2006, Larkum et al., 2007). Our work additionally demonstrates that bAP amplitude and Δ*Ca_AP_* can vary across branches, independent of distance from the soma. This variance can be explained by the branch structure of the dendritic tree. Previous studies have focused on attenuation of Δ*Ca_AP_* as a function of dendritic branch order (Spruston et al., 1995, Williams and Stuart, 2000); however, we find that explaining the observed variance in Δ*Ca_AP_* requires consideration of the full dendritic morphology, including the number of branches distal to the recording site and on sister dendrites. Elaborate branching patterns reduce input resistance, local bAP amplitude, and bAP-evoked calcium influx because dendritic input resistance is proportional to the surface area of membrane. We note that because bAPs are time varying signals, it is necessary to consider the complex impedance of each dendrite to fully explain bAP attenuation. As an alternative, we use computational models that account for complex impedance to demonstrate that bAP attenuation is predicted by the dendritic branching structure.

Our data cannot precisely resolve how much bAPs are attenuated in low Δ*Ca_AP_* branches because loose-patch recordings do not directly measure intracellular voltage (Figure 4) and voltage imaging signals are not precisely calibrated to absolute voltage (Figure 5). However, due to the differential impact on VGCC- and NMDAR-mediated calcium influx, we hypothesize that bAPs peak between −40 mV and −15 mV in low Δ*Ca_AP_* branches, based on our analysis in Figure 8. These results provide a key insight into the mechanisms underlying the intracellular heterogeneity of dendritic biophysical properties and bolsters our understanding of dendritic physiology.

### Activation of VGCCs in Low Δ*Ca_AP_* Branches

Our data indicate that bAP-dependent activation of VGCCs is branch-specific in cortical L2/3 pyramidal cells. Low Δ*Ca_AP_* branches exhibit little to no calcium influx evoked by single bAPs or even bursts of 5 bAPs at 150 Hz. In vivo measurements of L2/3 pyramidal cells have not reported high frequency bursts of bAPs (Smith et al., 2013, Wei et al., 2020), suggesting that in general, somatic depolarization does not lead to activation of VGCCs in low Δ*Ca_AP_* branches without additional depolarization from local synaptic input. We demonstrated that dendritic glutamate application can evoke VGCC-mediated calcium influx in low Δ*Ca_AP_* dendrites; however, this approach was designed to maximize local depolarization, rather than to resemble a physiological stimulus.

The lack of bAP-dependent activation of VGCCs in low Δ*Ca_AP_* branches may have important implications for neuronal function. First, because VGCC-mediated influx is responsible for multiple forms of synaptic and cellular plasticity (Kapur et al., 1998, Dolmetsch et al., 2001, Yasuda et al., 2003, Nevian and Sakmann, 2006, Scheuss et al., 2006, Brigidi et al., 2019), these data suggest that each of these forms of plasticity may be branch-specific in L2/3 pyramidal cells, at least with respect to their activation by bAPs. Second, these results have clear significance for the interpretation of dendritic calcium signals *in vivo.* A fundamental challenge in measuring dendritic activity with calcium imaging *in vivo* is distinguishing between calcium influx evoked by local synaptic input and global calcium signals, such as those evoked by bAPs (Xu et al., 2012, Beaulieu-Laroche et al., 2019, Francioni et al., 2019, Kerlin et al., 2019). Our results indicate that in low Δ*Ca_AP_* branches, bAPs do not evoke calcium signals without the addition of local synaptic input, which could make them a useful site for measuring local dendritic processing.

### Implications for Synaptic Plasticity

Previous work has demonstrated that bAP-evoked calcium influx selectively drives LTD while bAP-dependent amplification of synaptic calcium influx selectively drives LTP (Feldman, 2000, Matsuzaki et al., 2004, Nevian and Sakmann, 2006, Lee et al., 2009). Therefore, identifying branches with a selective reduction in bAP-evoked calcium influx while maintaining bAP-dependent amplification of synaptically-mediated calcium influx suggests that the ratio of LTD to LTP varies across branches in L2/3 pyramidal cells. Since the ratio of STDP-dependent depression to potentiation determines the cooperativity, strength, and sparsity of synaptic inputs (Song et al., 2000, Froemke et al., 2005, Sjostrom and Hausser, 2006), our work suggests that synaptic tuning properties may vary in a branch-specific manner based on the local dendritic branch structure. These findings complement a series of papers showing that calcium-dependent plasticity rules vary as a function of distance from the soma (Golding et al., 2002, Froemke et al., 2005, Sjostrom and Hausser, 2006, Gordon et al., 2006).

Our experiments were conducted in juvenile mice (between P21-P28). Although the dendritic branch structure is static after this age (Koleske, 2013, Richards et al., 2020), it is possible that changes in channel composition dampen the effects of dendritic branch structure on excitability. We show that such effects are possible with compartment models (Figure 7F), and it is known that changes in dendritic excitability can occur as a result of synaptic plasticity induction (Frick et al., 2003, Frick et al., 2004, Losonczy et al., 2008). Therefore, our findings may suggest a developmental function of low Δ*Ca_AP_* branches in shaping synaptic connectivity, consistent with the lower incidence of calcium influx in specific branches of L2/3 pyramidal cells during development in visual cortex (Yaeger et al., 2019).

### Implications for Dendritic Computation

Our work suggests that the dendritic branch structure of low Δ*Ca_AP_* branches can expand the representational capacity of neurons (Hausser and Mel, 2003). First, the lower input resistance of low Δ*Ca_AP_* branches makes them more compartmentalized from the rest of the cell, such that they may be more biased towards performing local computations (Polsky et al., 2004). Interestingly, our compartment models indicate that compartmentalization is asymmetric (Williams, 2004), such that forward propagation of synaptic input from low Δ*Ca_AP_* branches to the soma may be equivalent to high Δ*Ca_AP_* branches, despite their different biophysical properties. Second, branch-specific variation in calcium-dependent plasticity signals may diversify synaptic tuning, which would increase dendritic computational capacity (Poirazi et al., 2003, Hausser and Mel, 2003, Francioni and Harnett, 2021, Bicknell and Hausser, 2021). Together, these considerations indicate that low Δ*Ca_AP_* branches may be hotspots for dendritic computation due to their enhanced dependence on cooperative synaptic processing and higher likelihood of containing heterogeneous synaptic tuning.

## Material and Methods

### Genetic Constructs

For voltage imaging, we used QuasAr6a, an improved Archaerhodopsin-derived near-infrared genetically encoded voltage indicator (GEVI) (Tian et al., 2021). To improve expression and membrane trafficking we created the construct CAG::QuasAr6a-TS-dmCitrine-TSx3-ER2-p2a-jRGECO1a-CAAX, following the design in Ref. (Adam et al., 2019), where TS is the trafficking sequence from K_ir_2.1 (Gradinaru et al., 2010), dmCitrine is the non-fluorescent Y66G mutant of mCitrine (Adam et al., 2019), and ER2 is the endoplasmic reticulum export signal FCYENEV (Gradinaru et al., 2010). The self-cleaving p2a linker enabled bicistronic co-expression of a membrane-targeted Ca^2+^ indicator, jRGECO1a-CAAX. In the present studies, this indicator was only used for locating expressing neurons; for consistency with other experiments in this manuscript, we used the blue-shifted dye, Fluo-5F for Ca^2+^ imaging.

The genes were cloned into a second-generation lentiviral backbone with a CAG promoter (Addgene: #124775) using standard Gibson Assembly. Briefly, the vector was linearized by double digestion using BamHl and EcoRl (New England Biolabs, Ipswich, MA) and purified by the GeneJET gel extraction kit (ThermoFisher, Waltham, MA). DNA fragments were generated by PCR amplification and then fused with the backbones using NEBuilder HiFi DNA assembly kit (New England Biolabs). All plasmids were verified by sequencing (GeneWiz, Cambridge, MA).

### *In utero* electroporation (IUE)

All procedures involving animals were in accordance with the National Institutes of Health Guide for the care and use of laboratory animals and were approved by the Harvard University Institutional Animal Care and Use Committee (IACUC). The IUE surgery was made as described previously (Kwon and Sabatini, 2011). Embryonic day 15.5 (E15.5) timed-pregnant female CD1 mice (Charles River, Wilmington, MA) were deeply anesthetized and maintained with 2% isoflurane. The animal body temperature was maintained at 37 °C. Uterine horns were carefully exposed, and periodically rinsed with warm PBS. The plasmid DNA was diluted with PBS (2 μg/μL; 0.005% fast green), and 1 μL of the mixture was injected into the left lateral ventricle of pups. Electric pulses (40 V, 50 ms duration) were delivered five times at 1 Hz using a tweezers electroporation electrode (CUY650P5; Nepa Gene, Ichikawa, Japan). Injected embryos were placed back into the abdominal cavity, and the surgical wound was sutured with PGCL25 absorbable sutures (Patterson, Saint Paul, MN).

### Slice Preparation

Acute coronal slices were prepared from young adult, C57Bl/6j wild-type mice (Jackson Labs, Bar Harbor, ME) between postnatal days P21-P28. Animals were anesthetized via inhalation of isoflurane then immediately decapitated. The brain was rapidly removed and placed in ice-cold cutting artificial cerebrospinal fluid (ACSF) containing (in mM): 125 NaCl, 2.5 KCl, 25 NaHCO_3_, 1.25 NaH_2_PO_4_, 1 CaCl_2_, 10 MgCl_2_, and 25 glucose, saturated with 95% O_2_ and 5% CO_2_. We cut coronal slices with a VT1200S Vibratome (Leica, Buffalo Grove, IL) while maintained in the cutting solution. Slices were then incubated at 34°C for 30 minutes in a recovery ACSF containing (in mM): 92 NaCl, 28.5 NaHCO_3_, 2.5 KCl, 1.25 HaH_2_PO_4_, 2 CaCl_2_, 4 MgCl_2_, 25 glucose, 20 HEPES, 3 Sodium Pyruvate, and 5 Sodium Ascorbate, saturated with 95% O_2_ and 5% CO_2_. Slices were next transferred to a 20°C room in recovery ACSF until use (>30 minutes after slicing) and not held for longer than 8 hours. Experiments were conducted at 34°C in recording ACSF containing (in mM): 125 NaCl, 2.5 KCl, 25 NaHCO_3_, 1.25 NaH_2_PO_4_, 1.5 CaCl_2_, 1 MgCl_2_, and 25 glucose, saturated with 95% O_2_ and 5% CO_2_. All solutions were prepared with osmolality between 300-310 mOsm/kg, adjusted with either water or glucose. For voltage imaging experiments in Figure 5, coronal slices were prepared from CD1 mice between P21-P28. The slicing solution contained (in mM): 210 sucrose, 3 KCl, 26 NaHCO_3_, 1.25 NaH_2_PO_4_, 5 MgCl_2_, 10 D-glucose, 3 sodium ascorbate and 0.5 CaCl_2_, and was saturated with 95% O_2_ and 5% CO_2_. The slices were transferred to an incubation chamber containing the recording solution (in mM): 124 NaCl, 3 KCl, 26 NaHCO_3_, 1.25 NaH_2_PO_4_, 2 MgCl_2_, 15 D-glucose and 2 CaCl_2_, and was saturated with 95% O_2_ and 5% CO_2_.

### Electrophysiology

Somatic whole-cell recordings were acquired from cortical excitatory L2/3 pyramidal cells using IR-DIC. Patch pipettes (2-4.5 MΩ) were filled with an internal solution containing (in mM): 130 K-Gluconate, 10 KCl, 10 HEPES, 4 MgATP, 0.5 Na_2_GTP, 10 Phosphocreatine-disodium salt, 0.3 Fluo-5F and 0.01 Alexa 594. For voltage-imaging experiments in Figure 5, the internal solution contained (in mM): 8 NaCl, 130 KMeSO_3_, 10 HEPES, 5 KCl, 0.5 EGTA, 4 Mg-ATP, 0.3 Na3-GTP, 0.3 Fluo-5F, and 0.01 JF549i. The pH was adjusted to 7.3 using KOH and osmolality was adjusted to 285-295 mOsm/Kg with water. In a subset of experiments shown in Figure 2E, 300 μM Fluo-4 was used instead of Fluo-5F, but amplitudes of calcium signals were comparable, so we merged the data. All recordings were performed with a Multiclamp 700B amplifier (Molecular Devices, San Jose, CA). Pipette capacitance was neutralized prior to break-in and the series resistance was fully balanced for all recordings. Series resistance ranged from 7-25 MΩ. We elicited APs by injecting a 1-2 ms current of 0.5-3.5 nA. Recordings were aborted if the cell’s membrane potential exceeded −60 mV, if the dendrites begun filling with calcium (as indicated by the baseline Fluo-5F signal), or if the series resistance became too high for us to reliably evoke action potentials. For experiments in which we performed glutamate uncaging, we also aborted recordings if the input resistance of the cell exceeded 200 MΩ. When we performed glutamate uncaging, we added the following drugs via bath application: 3.75 mM MNI-Glutamate (for uncaging), 10 μM 2-CA (to reduce presynaptic release probability and maintain quiescent recording conditions), 1 unit/mL glutamate pyruvate transaminase (which catalyzes free glutamate into α-ketoglutarate), and 3 mM Sodium Pyruvate (a necessary cofactor for GPT), to the recording ACSF. In experiments where we blocked uncaging-evoked depolarization in Figure 4, we added 10 μM NBQX to the recording ACSF.

Dendritic loose-patch recordings were acquired from cortical excitatory L2/3 pyramidal cell dendrites after acquiring somatic whole-cell recordings and filling the cells with 10 μM Alexa 594. The open-tip resistance of dendritic pipettes was 8-12 MΩ, and the pipettes were filled with recording ACSF. Dendritic recordings were made with a Multiclamp 700B amplifier (Molecular Devices) in current-clamp mode. We used scanning DIC and two-photon imaging to target pipettes to dendritic branches. Before acquiring a loose-patch seal, we pushed the dendritic pipette into the dendrite until we observed a visible kink in the dendrite. For each dendritic recording, we recorded the electrical signal evoked by an AP before and after applying a brief pulse of negative pressure to achieve a loose-patch seal (Figure 4 – figure supplement 1). All dendritic analyses in Figure 4 are derived from the difference between these two signals, which removes small electrical artifacts of somatic current injection and focuses our analysis on the signal arriving directly from the dendritic membrane.

The glutamate puff solution used in Figure 3 was applied with a large patch-pipette (<1 MΩ) connected to a picospritzer. The puff solution was composed of recording ACSF in addition to 1 mM glutamate, 20 μM CPP and 50 μM MK-801. For glutamate puff experiments, we also added 20 μM CPP and 50 μM MK-801 to the recording ACSF to block NMDARs. In some experiments, we puffed recording ACSF onto the dendrite without glutamate or NMDAR blockers to control for displacement artifacts evoked by the positive pressure (Figure 3 – figure supplement 1). We used scanning DIC and two-photon imaging to bring puff pipettes near dendrites and used a short 5-15 ms puff to apply glutamate to dendritic branches. The pressure of the puff was increased gradually between 5-20 PSI until a clear somatic depolarization and dendritic calcium signal were observed.

### Two-photon Imaging and Glutamate Uncaging

Two-photon imaging was performed with a custom-built microscope described previously (Carter and Sabatini, 2004). We tuned a mode-locked femtosecond laser to 840 nm to excite both Fluo-5F and Alexa 594. All calcium imaging was performed >10 minutes after break-in to allow fluorophores to passively diffuse into the dendritic tree. Stimulus-evoked changes in fluorescence were quantified as the change in green (Fluo-5F) fluorescence relative to a baseline period, divided by the baseline red (Alexa 594) fluorescence (ΔG/R) (Bloodgood and Sabatini, 2007). We used a custom algorithm to center the field of view on each trial (Carter and Sabatini, 2004). We measured fluorescence with a 2 ms line-scan through the recording sites (at least 1 dendritic shaft and 0-3 dendritic spines depending on the experiment) and averaged over the spatial coordinates of each compartment to extract a timevarying trace (Figure 1). We waited at least 10 s in between trials to minimize photodamage and allow the calcium signal to fully return to baseline. In the experiments in Figure 6 – figure supplement 2, we performed point-scans on dendritic spines to maximize the sampling rate (125 kHz and downsampled to 8 kHz to minimize shot noise).

To perform laser photolysis of MNI-glutamate, we used a short, 500 μs femtosecond laser pulse tuned to 725 nm. For each experiment, we gradually increased laser intensity until either a 0.5 mV EPSP was generated or a clear calcium signal in the dendritic spine was evoked. We moved the uncaging spot around the dendritic spine to maximize the EPSP size as an attempt to uncage glutamate directly apposed to the postsynaptic density.

### Simultaneous voltage and calcium imaging

Interleaved voltage- and calcium-imaging experiments in Figure 5 were conducted on modified homebuilt structured illumination epifluorescence microscope previously reported (Adam et al., 2019). Briefly, blue illumination was patterned by a digital micromirror device (DMD) and used for structured illumination calcium imaging via the calcium-sensitive dye Fluo-5F. Yellow illumination was also patterned by a DMD and used for structured illumination mapping of dendritic morphology via a dye JF549i. Red illumination was patterned by a holographic spatial light modulator (SLM) and used for structured illumination voltage imaging.

Laser lines from a blue laser (488 nm, 150 mW, Obis LS) and green laser (561 nm, 150 mW, Obis LS) were combined, sent through an acousto-optic modulator for amplitude control, and expanded onto a DMD (V-7000 VIS, ViALUX) for spatial modulation. The DMD was then re-imaged onto the sample via a 25x water-immersion objective, NA 1.05, Olympus XLPLN-MP. A red laser (MLL-FN-639, 1 W, CNI lasers) was expanded onto an SLM (P1920-0635-HSP8, Meadowlark, Frederick, CO), combined with the blue and yellow lasers via a polarizing beamsplitter, and re-imaged onto the back aperture of the objective. Fluorescence was collected by the objective and imaged onto a sCMOS camera (Hamamatsu Orca Flash 4.0) with the appropriate emission filter for each indicator. Voltage-imaging recordings were acquired at a 1 kHz frame rate. Calcium imaging was recorded at 20 Hz.Two-photon imaging and reconstruction was performed with a custom-built microscope adapted to be combined with the 1P illumination. Maximum intensity projections of z-stacks were used to form images of the dendritic arbor.

### Physiology and Imaging Analysis

All analysis was performed using custom software written in MATLAB and Python. All plots with error bars show mean +/- standard error. To compute the amplitude and peak time of uEPSPs, we first averaged across trials, subtracted the baseline voltage, and applied a median filter with a 1 ms window. Then, we found the maximum voltage between 0-25 ms after uncaging and defined this as the amplitude and peak time. The amplitude of stimulus-evoked calcium signals (ΔG/R) was computed using a 10 ms average of the fluorescence signal centered around the average peak time for each recording site. To compute the amplitude of dendritic electrical signals in Figure 4, we first computed the average electrical signal in a 0.2 ms window surrounding the peak voltage for each trial. The amplitude of each recording was defined as the average peak voltage from the trials in the top 50^th^ percentile signal-to-noise ratio (peak divided by baseline standard deviation). While this method is unsatisfactory for proper comparisons of amplitude across recordings, we only use these data to determine if somatic APs evoke a measurable electrical signal in the dendrites. For measurements of the ratio of Δ*Ca_AP_* from within segment or across branch point pairs (Figure 6 – figure supplement 1A-C), we only considered pairs if the proximal site had a Δ*Ca_AP_ >* 0.1 because the ratio of Δ*Ca_AP−dist_/ΔCa_AP−prox_* is not well-defined if the proximal site failed to exhibit AP-evoked calcium influx. Δ*Ca_AP_* never dropped to ~0 and then recovered in more distal sites along the same branch.

For analysis of epifluorescence voltage- and calcium-imaging data in Figure 5, we created a spike-triggered average (STA) movie comprising the average bAP-evoked signal. An initial estimate of the STA waveform for each dendritic branch was calculated by averaging the pixel values in a region of interest comprising the 5% of pixels with greatest mean brightness. Maps of the baseline (F) and spike-dependent (Δ*F*) signals were then calculated pixel-by-pixel by linear regression of the STA movie against a normalized copy of the initial STA waveform estimate, after accounting for a pixel-dependent offset. To account for spatially variable background, watershed segmentation was applied to the Δ*F* map to identify regions with high spike-dependent signal. For the region surrounding pixels with the highest ΔF, a pixel-by-pixel plot of *F* vs. Δ*F* was fit to a line via linear regression. The inverse of the slope was the calculated Δ*F/F_0_*

### Morphological Analysis

After all experiments, we collected two-photon z-stacks of each cell to map the dendritic morphology between the soma and the recording sites. In the neurons used in Figure 6, we imaged the entire apical dendritic tree after finishing experiments to reconstruct the cells for morphological analyses and compartmental modeling in NEURON. For every recording site, we measured the on-path distance from the soma using maximum z-stack projections of each cell. The distance for each recording site is a slight underestimate of the true on-path distance from the soma because we ignored changes along the z-axis. All imaging was conducted close to the surface of the slice and the maximal angle between the soma and any recording site was 25°, so errors in measured distance were <10%.

We developed a metric for branch impedance that fully accounts for the extent of nearby dendritic branching, independent of the structure of the dendritic tree. First, we defined the term *β* as the number of dendritic branches at a given on-path distance from a recording site (Figure 6G). *β* always starts as 2 (one on each side of the recording site) and increases by 1 after every branch point. This is distinct from Scholl analysis as it measures distance based on the on-path distance rather than absolute distance. Next, we performed point-wise multiplication between *β* and an exponential filter to discount for dendritic branches that are further away from the recording site. Finally, we computed branch impedance as 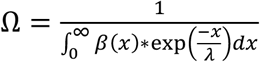, which is smallest at sites that are closest to large numbers of dendritic branches and is theoretically proportional to electrotonic input resistance.

The true electrotonic input resistance is analytically determined by the cable length constant, which is not feasible to measure in thin caliber L2/3 pyramidal cell dendrites. As an alternative, we estimated how much membrane is simultaneously depolarized by a bAP as an estimate of how much of the dendritic tree can affect the depolarization at a particular site. We used a rapid point-scan imaging approach that allowed us to measure Δ*Ca_AP_* in dendritic spines at >8 kHz (Figure 6 – figure supplement 2A). We plotted the latency to peak influx (measured from the derivative of fluorescence) as a function of the distance from the soma and used linear regression to estimate the conduction velocity (Figure 6 – figure supplement 2A-B). We found that the conduction velocity of bAPs in L2/3 dendrites is 154.9 μm/ms. In the same cells used to measure conduction velocity, we plotted the somatic AP waveforms and measured their full-width half-max (Figure 6 – figure supplement 2C-D). The somatic AP has a duration of about 0.94 ms (Figure 6 – figure supplement 2C-D). To compute the spatial width of a propagating bAP, we multiplied the conduction velocity by the duration (154.9 μm/ms * 0.94 ms = 145.6 μm). This estimate of bAP width led to a choice of length constant that maximized the correlation between branch impedance and input resistance (Figure 6 & Figure 6 – figure supplement 3).

### Biophysical Compartmental Modeling

Simulations used in Figure 6F, Figure 7, and Figure 7 – figure supplement 1 were performed in the NEURON simulation environment (Carnevale and Hines, 2006) using Python (Version 3.7). We reconstructed all 8 neurons used in Figure 6 with maximum z-projections of the Alexa 594 fluorescence signal. Each compartmental model was composed of a cylindrical soma with length and diameter equal to 12 μm, a 30 μm axon initial segment (AIS) with a 2 μm diameter connected to the soma, a 300 μm axon with a 1 μm diameter connected to the AIS, and a dendritic tree connected to the soma that with identical branch structure to the 8 neurons used in Figure 6 and a uniform diameter of 1 μm. We explored distance-dependent decreases in diameter but found qualitatively similar results so did not show these experiments. In some experiments, we performed simulated cuts of the dendritic tree in NEURON. For each cell, we removed branches close to the low Δ*Ca_AP_* recording site until the input resistance of the high and low Δ*Ca_AP_* sites were comparable.

We used standard passive parameters: C_m_ = 1 μF/cm^2^, Rm=7000 Ω-cm^2^, Ri = 100 Ω-cm. Active conductances were based on a previous model of L2/3 pyramidal cells that was fit to in vivo somatic and dendritic recordings (Smith et al., 2013). Dendritic active conductances were completely uniform, such that all differences across dendrites were determined by the distance from the soma and the dendritic branch structure. Each model contained the following channels with given conductance density in pS/μm^2^: voltage-gated sodium channels (axon: 170, AIS: 2550 soma: 85, dendrites: 85), voltage-gated potassium channels (axon: 33, AIS: 100, soma: 100, dendrites: 3), M-Type potassium channels (soma: 2.2, dendrites: 1), calcium-dependent potassium channels (soma: 3, dendrites: 3), high voltage-activated calcium channels (soma: 0.5, dendrites: 0.5), and low voltage-activated calcium channels (soma: 3, dendrites: 1.5). Additionally, we added an A-Type potassium channel (Migliore et al., 1999) to the soma and dendrites with a variable conductance density that was set for each cell independently. For experiments in Figure 6F, Figure 7A-E, Figure 7G-H, and Figure 7 – figure supplement 1, we used a Nelder-Mead simplex algorithm (from the SciPy toolbox in Python) to set the A-Type potassium channel conductance density such that backpropagating APs peaked at −10 mV in the high Δ*Ca_AP_* recording site within each pair (values for each site plotted in left-most column of points in Figure 7F). The computed density was used for the soma and entire dendritic tree. We chose to set the AP peak at −10 mV because it is consistent with the AP amplitude measured with whole-cell recordings from L2/3 pyramidal cell dendrites >100 μm from the soma (Waters et al., 2003, Larkum et al., 2007, Smith et al., 2013). The same values for A-Type potassium channel density were used in experiments with simulated cut dendrites. The values shown in Figure 7F use the same algorithm to select A-Type potassium channel density for high and low Δ*Ca_AP_* recording sites independently, for both the intact and cut dendritic branch structure.

### Conductance Simulations

We computed the voltage-dependent open probability and time constant of VGCCs and NMDARs using published data (Jahr and Stevens, 1990, Reuveni et al., 1993). For VGCCs, we used the following voltage-dependent forward and backward rate constants (*α_v_* and *β_v_*, respectively) of the activation and inactivation gates (in mV):

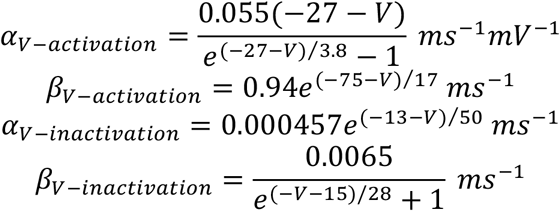

To compute open probability and time constant for the activation (denoted ‘*m*’) and inactivation (denoted ‘*h*’) gates of VGCCs, we used the following equations: *P_open_*(*V*) = *α_v_*/(*α_v_* + *β_v_*), *τ*(*V*) = 1/(*α_v_* + *β_v_*). Because the time constant of the inactivation gate is so slow (the minimum time constant is 170 ms between −80 and +40 mV), it acts as a constant, so we did not plot it in Figure 8B, although it was used in the simulations in Figure 8C-E.

For NMDARs (denoted ‘*n*’), we calculated the voltage-dependent open-probability and time constant as *P_open_* (*V*) = *k_off_*/(*k_on_* + *k_off_*) and *τ*(*V*) = 1/(*k_on_* + *k_off_*) based on the measured on and off rates of the Mg^2+^ block (Jahr and Stevens, 1990), with [Mg^2+^] = 1 μM:

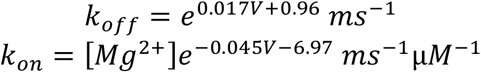

For numerical integration in Figure 8C, we used Euler’s method to compute changes in the state of each gate with Δ*t* = 0.01 *ms.* We used the following differential equations:

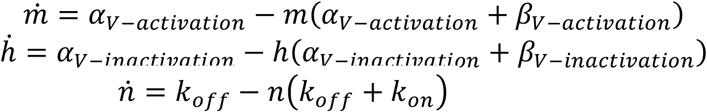

The open probability of NMDARs is equal to *n,* and the open probability of VGCCs is equal to *m^2^h.* To convert the open probability into a voltage-dependent calcium current, we used a modified version of the Goldman-Hodgkin-Katz current equation with [*Ca*]_*in*_ = 75 *nM* and [*Ca*]_*out*_ = 1.5 *mM:*

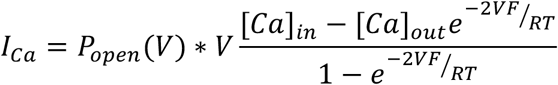

We used quadratic voltage depolarizations to mimic the shape of APs. For each stimulus, we used a preset amplitude (*V_amp_*) and duration (*V_dur_*). Then, we solved the following equation for *a*: 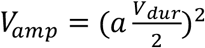, such that the stimulus would have a maximum height of *V_amp_* along the domain 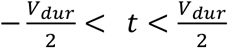. Then, we added the equation: *step*(*t*) = *V_amp_* – (*at*)^2^ to the baseline voltage of −70 mV.

To determine the integrated calcium influx in Figure 8E, we integrated the calcium current evoked by the voltage step after subtracting the baseline current (NMDARs are partially open at rest).

## Acknowledgements

We thank all members of the Sabatini Lab for comments and advice, in particular R. Hakim, E. Sayed, P. Capelli, S. Melzer, K. Reinhold, T. Kula, A. Granger, and S. Kim for suggestions on the manuscript. We also thank J. Reggiani, N. Hahn, W. Regehr, C. Harvey, and B. Bean for helpful suggestions and critical insights. This work was supported by 5F31NS113353-03 from NINDS to ATL, R37NS046579 from NINDS to BLS, NIH R01 1RF1MH117042-01 to AEC, a Vannevar Bush Faculty Fellowship and a Brain Research Foundation Scientific Innovation Award to AEC, and a grant from the Harvard Brain Initiative to BLS and AEC. JDW is a Merck Awardee of the Life Sciences Research Foundation.

---

**Figure 1 – figure supplement 1:**
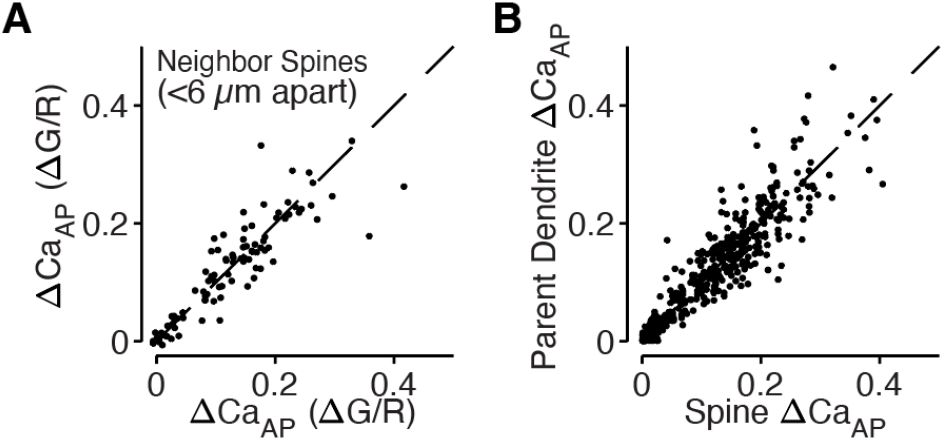
bAP-evoked calcium influx is regulated across dendritic branches and consistent in neighboring dendritic spines. A) Comparison of ΔCa_AP_ in neighboring dendritic spines measured simultaneously. The strong correlation indicates that nearby spines have similar calcium signals throughout the dendritic tree and intracellular differences in ΔCa_AP_ are only observed across branches. B) Comparison of ΔCa_AP_ between dendritic spines and their parent dendritic shafts. The strong correlation indicates that ΔCa_AP_ is regulated at the level of dendritic branches and motivates our use of dendritic calcium influx as a proxy for nearby dendritic spines.

**Figure 2 – figure supplement 1:**
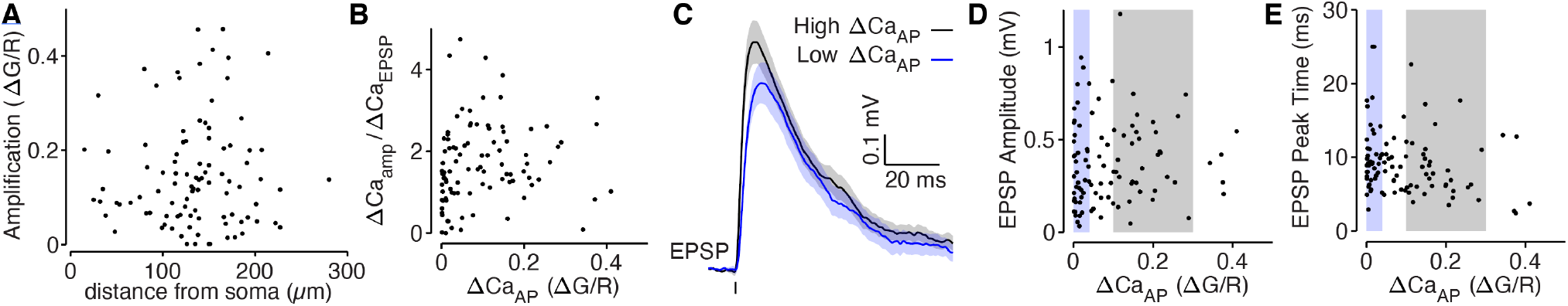
bAP-dependent amplification is not determined by glutamate uncaging properties. A) bAP-dependent ΔCa_AP_ of uEPSP-evoked calcium influx across distance from soma in all dendritic spines. B) Comparison of bAP-dependent amplification divided by calcium evoked by the EPSP (to normalize for the level of NMDAR activation) with bAP-evoked calcium influx. The plot is qualitatively similar to the plot in Figure 2B, suggesting that differences in NMDAR activation do not explain the results. C) Average EPSP recorded from the soma for high and low ΔCa_AP_ dendritic spines. D) Comparison of EPSP amplitude with bAP-evoked calcium influx. E) Comparison of EPSP peak time with bAP-evoked calcium influx. The small negative correlation is primarily because low ΔCa_AP_ dendritic spines tend to be further from the soma, so the slower EPSPs are primarily due to cable filtering rather than synaptic properties.

**Figure 3 – figure supplement 1:**
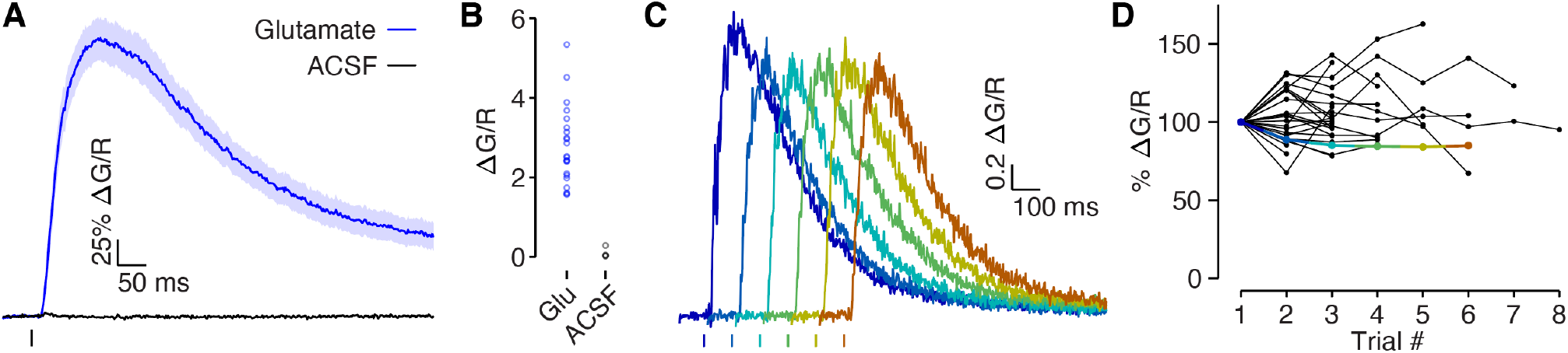
Glutamate puff-evoked calcium signals are mediated by voltage-gated calcium channels. A) Dendritic calcium influx evoked by picospritzer puff of either 1 mM glutamate (N=22) or ACSF (N=4). B) Comparison of peak calcium influx evoked by 1 mM glutamate and ACSF. No calcium influx was observed in response to ACSF, indicating that the calcium signal was not due to mechanical displacement evoked by the picospritzer. C) Example traces of calcium influx evoked by 1 mM glutamate puff for one dendrite across all trials, shifted along the x-axis to visualize each trial clearly. D) Dendritic calcium influx evoked by 1 mM glutamate for each trial, normalized to first trial amplitude. If glutamate was out-competing CPP (a competitive antagonist) at NMDAR’s binding sites, then the presence of MK-801, a use-dependent blocker, should reduce NMDAR conductance in each trial. No systematic reduction in the calcium signal was observed, indicating that NMDARs were fully blocked. Colored points indicate example used in panel C.

**Figure 4 – figure supplement 1:**
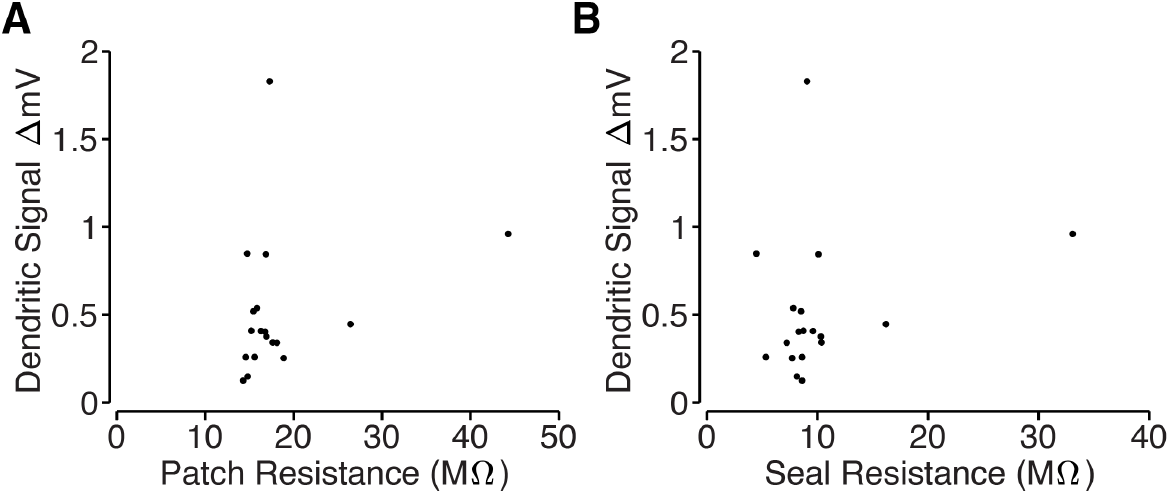
Loose-patch recording properties. A) Comparison of total resistance of loose-patch recordings and amplitude of the dendritic electrical signals. The total resistance includes the pipette resistance and the seal resistance, which was measured after acquiring a loose-patch seal on a target dendrite using a small current step. B) Comparison of the seal resistance and the amplitude of the dendritic electrical signals. Seal resistance was computed by subtracting the loose-patch resistance (measured after acquiring a loose-patch seal) from the pipette resistance (measured with the pipette in the bath solution). Note: several unmeasured noise sources prohibit inference of intracellular bAP amplitude based on loose-patch recordings. We are primarily measuring capacitive currents, 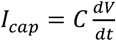, but we cannot directly measure dendritic capacitance and may differ across dendritic branches. Also, the amplitude of the recording is also proportional to the true seal resistance (Figure 4D); however, our loose-patch recordings used large pipettes (pipette resistance ≈ 8 MΩ) and the total patch resistance only increased by ~10 MΩ (Figure 4C), so estimates of the seal resistance are likely to be contaminated by the neuropil.

**Figure 6 – figure supplement 1:**
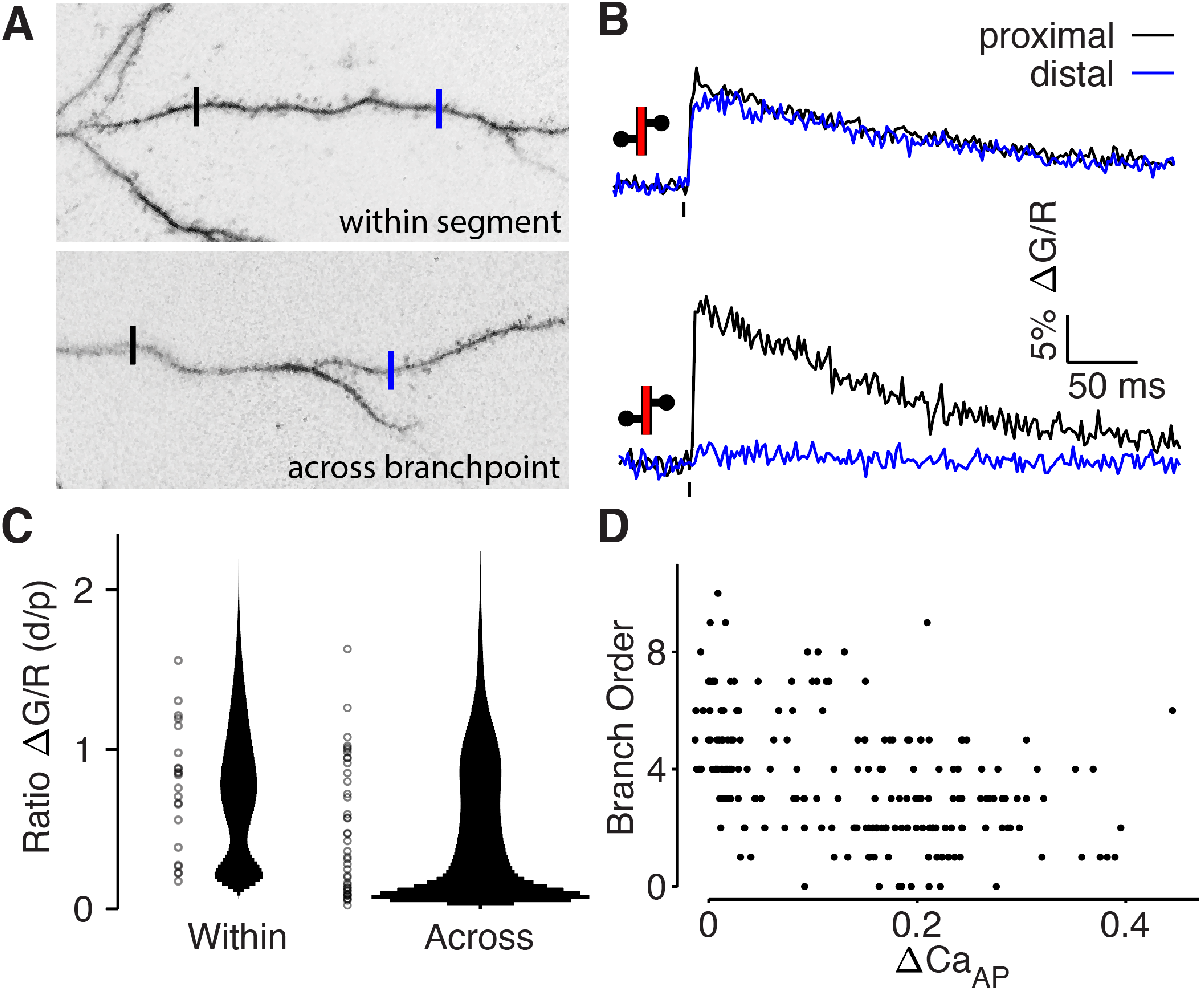
bAP-evoked calcium influx attenuates around branch points. A) Alexa 594 MZP showing two ROIs within a segment of dendrite or across a branch point. Blue ROI is more distal than black ROI. B) Calcium influx evoked by bAP in ROIs from panel A. C) Distribution of the ratio of peak calcium influx (distal/proximal) for within-segment pairs and across branchpoint pairs. Width of bar indicates number of observations. Rank-sum: p=0.0145. D) Comparison of dendritic bAP-evoked calcium influx and branch order.

**Figure 6 – figure supplement 2:**
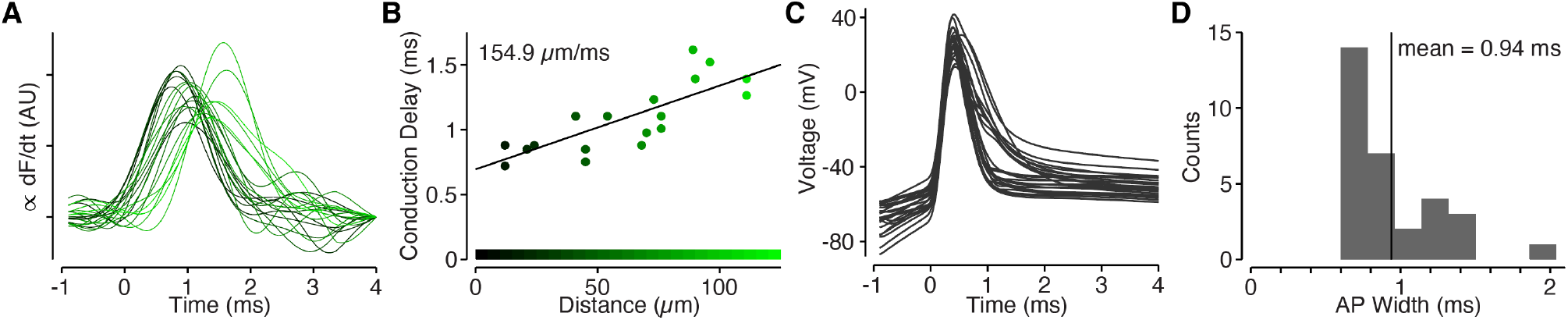
Back-propagating APs span 145 μm in L2/3 cell dendrites. A) Derivative of ΔCa_AP_ in dendritic spines, color-coded by their distance from the soma (color scale in panel B). We used a point-scan imaging approach in which we sampled the spines calcium-dependent fluorescence signal at 8 kHz to accurately measure the time-course of calcium influx in each spine. B Conduction delay of ΔCa_AP_ (measured at peak of derivative) as a function of distance. The inverse of the slope gives the conduction velocity of APs as they back-propagate through L2/3 cell dendrites. C) Waveforms of APs recorded from L2/3 cells aligned to peak of second derivative (kink of AP waveform). D) Histogram of FWHM of AP waveform for each cell. To compute the spatial spread of APs in L2/3 cell dendrites, we multiplied the average AP width by the conduction velocity 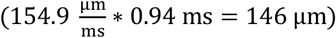.

**Figure 6 – figure supplement 3:**
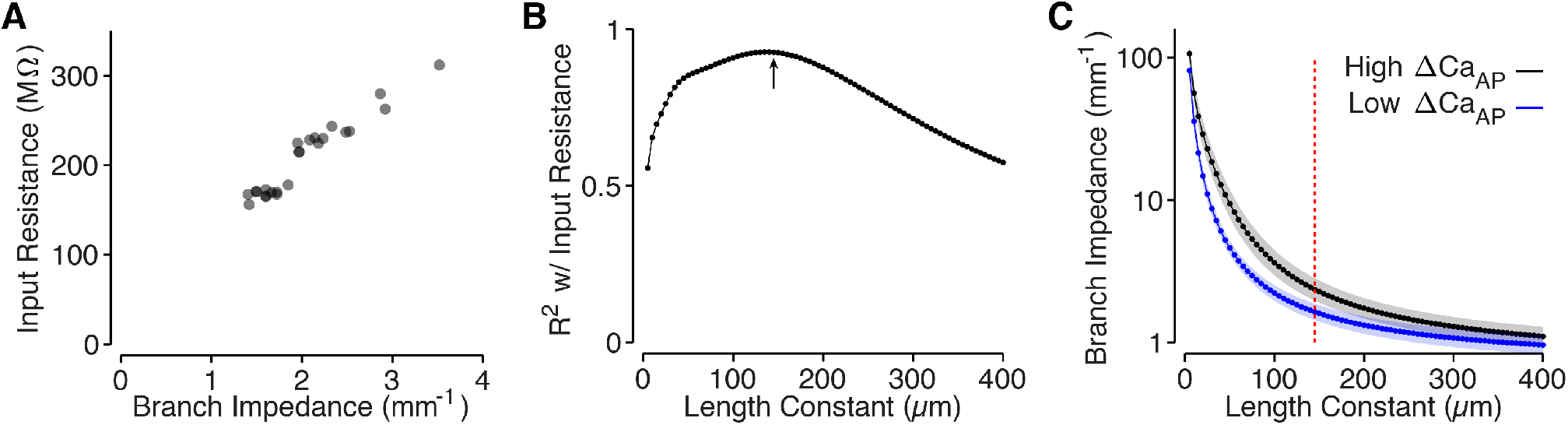
Branch impedance is correlated with input resistance. A) Input resistance of each site, measured in NEURON, compared to the branch impedance of each site, measured based on the cell’s dendritic morphology using a length constant of 145 μm. R^2^=0.93 at λ=145. B) Correlation (R^2^) between branch impedance and input resistance for all length constants of branch impedance. The arrow indicates the length constant used in panel A and Figure 6, computed based on the shape of the AP in L2/3 cell dendrites. C) Comparison of branch impedance between high and low ΔCa_AP_ dendrites for each length constant of branch impedance. Red dotted line indicates length constant used in panel A and Figure 6. Y-axis is on a log scale.

**Figure 7 – figure supplement 1:**
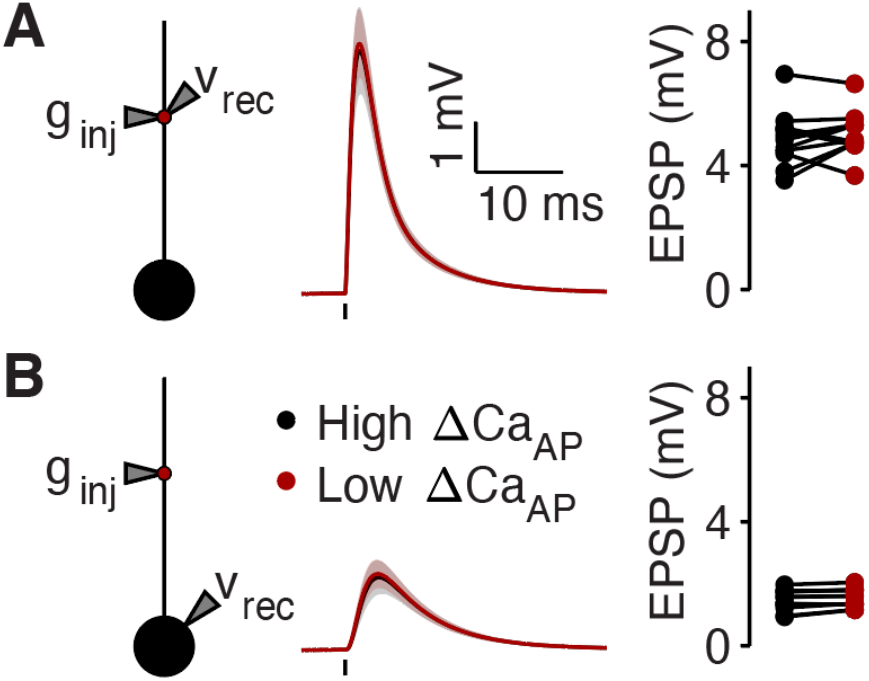
Cutting dendrites normalizes local EPSP amplitude. A) Same as in Figure 7G, but with cut dendrites. Left: a synaptic conductance was injected into each site and the local EPSP was recorded in NEURON compartment models with cut dendrites. Middle: dendritic EPSPs recorded in high (black) and low (maroon) ΔCa_AP_ dendritic sites. Right: comparison of dendritic EPSP amplitudes recorded in each high/low ΔCa_AP_ pair. B) Same as in Figure 7H, but with cut dendrites. Synaptic conductance injected into dendrite and recorded in the soma.

